# Characterization and therapeutic suppression of KEAP1-NRF2-driven resistance to KRAS inhibitors in pancreatic and lung cancer

**DOI:** 10.64898/2026.04.18.719329

**Authors:** Wen-Hsuan Chang, Alec J. Vaughan, Addison G. Stamey, Mariana Mancini, Makiko Hayashi, Runying Yang, Ryan Robb, Daniel Andrussier, Jeffrey A. Klomp, Andrew M. Waters, Antje Schaefer, Brian M. Wolpin, Kirsten L. Bryant, Adrienne D. Cox, Fernando Moreira Simabuco, Kwok-Kin Wong, Andrew J. Aguirre, Clint A. Stalnecker, Thales Papagiannakopoulos, Channing J. Der

**Affiliations:** Lineberger Comprehensive Cancer Center, University of North Carolina at Chapel Hill, Chapel Hill, North Carolina; Department of Pathology, NYU Langone Health, New York, New York; Laura and Isaac Perlmutter Cancer Center, New York University Langone Health, New York, New York; Department of Biochemistry, Federal University of São Paulo, São Paulo, Brazil; Department of Pharmacology, University of North Carolina at Chapel Hill, Chapel Hill, North Carolina; Department of Medical Oncology, Dana-Farber Cancer Institute, Boston, Massachusetts; Department of Radiation Oncology, University of North Carolina at Chapel Hill, Chapel Hill, North Carolina; The Broad Institute of Harvard and MIT, Cambridge, Massachusetts, USA; Harvard Medical School, Boston, Massachusetts; Department of Medicine, Brigham and Women’s Hospital, Boston, Massachusetts; Department of Biochemistry and Biophysics, University of North Carolina at Chapel Hill, Chapel Hill, North Carolina

## Abstract

The recent approval of KRAS inhibitors supports the therapeutic value of targeting mutant KRAS cancers. However, clinical efficacy is hindered by both primary and treatment-associated acquired resistance. We applied a CRISPR-Cas9 loss-of-function screen and identified loss of *KEAP1* as a resistance mechanism to the KRAS^G12D^-selective inhibitor MRTX1133 and the RAS(ON) multi-selective inhibitor RMC-7977 in pancreatic cancer models. RNA-sequencing analyses revealed a *KEAP1*^KO^ transcriptome that is distinct from the ERK-, MYC-, and YAP/TAZ-TEAD-dependent transcriptional programs that drive KRAS inhibitor resistance, demonstrating a distinct mechanism of resistance. We then established a PDAC KEAP1-deficient (PKD) gene signature that was enriched in patients and preclinical models insensitive to KRAS inhibitor treatment. Finally, we observed that KEAP1-deficient cells exhibited elevated glutamine metabolism, and combination treatment with the glutamine antagonist DRP-104 (sirpiglenastat) enhanced KRAS inhibitor suppression of pancreatic and lung tumors.

**SIGNIFICANCE:** KEAP1 loss is associated with reduced response to KRAS inhibitor therapy. We demonstrate that *KEAP1* loss-associated resistance can be overcome by pharmacologic inhibition of the *KEAP1* loss-induced glutamine dependency, establishing a combination to enhance RAS inhibitor clinical efficacy.

## INTRODUCTION

Mutational activation of *KRAS* is associated with ∼20% of all cancers, with the highest frequencies found in pancreatic ductal adenocarcinoma (95%; PDAC), and prevalent in lung adenocarcinoma (37%; LUAD) (1,2). Although KRAS was once considered undruggable (3), the recent approval of two KRAS inhibitors (sotorasib and adagrasib) selective for the glycine-12-cysteine (G12C) smoking associated mutation for KRAS^G12C^-mutant non-small cell lung cancer (NSCLC) and colorectal cancer (CRC) has shattered that perception (4–7).

Following the successful development of KRAS^G12C^-selective inhibitors, many inhibitors have been developed and entered clinical evaluation that target essentially all KRAS mutant alleles (3,8). Similar to the approved KRAS^G12C^ inhibitors, many are mutation-selective, especially for the most prevalent KRAS mutation in cancer, KRAS^G12D^ (9,10). Moreover, the first-in-class RAS(ON) multi-selective inhibitor daraxonrasib (RMC-6236) has shown excellent activity as second-line treatment in KRAS-mutant PDAC, comparable to responses to first-line chemotherapy but without the severe toxicity seen with standard-of-care cytotoxic drugs (11,12).

Despite the impressive clinical activities seen with KRAS inhibitors, depth and durability of response limit their long-term efficacy (3,13). Clinical evaluation of KRAS^G12C^ inhibitors in *KRAS*-mutant NSCLC, CRC, and PDAC patients have reported median overall response rates (ORR) generally below 50% with the duration of response (DOR) typically under 12 months, underscoring primary/innate resistance and the rapid emergence of acquired resistance, respectively (3). Acquired resistance also limits the long-term efficacy of daraxonrasib (14,15). Thus, a better understanding of the mechanisms underlying resistance to KRAS inhibitors is essential for developing more effective combination therapeutic strategies.

Evaluation of DNA alterations identified putative mechanisms of acquired resistance in only ∼50% of patients who relapsed on KRAS^G12C^ and pan-RAS inhibitors, and most (>90%) mutations are in components in the RAS signaling network. Non-genetic transcriptional mechanisms have also been identified in preclinical studies (3). For example, prolonged treatment with KRAS inhibitors was shown to activate compensatory pathways, including TEAD and MYC transcription factors as well as the RAF-MEK-ERK signaling network, thereby promoting tumor survival and drug resistance (16–22).

The dysregulation of the KEAP1-NRF2 pathway has emerged to be a major factor limiting the efficacy of KRAS^G12C^ inhibitors (14,16,17,23). KEAP1, a component of Cullin 3-E3 ubiquitin ligase complex, regulates the expression of the NRF2 transcription factor by targeting it for ubiquitin-dependent proteasomal degradation (24–26). Under oxidative stress, KEAP1 undergoes conformational changes, leading to NRF2 stabilization and subsequent transcriptional activation of antioxidant and cell survival genes. In cancer cells, oncogenic mutations on KEAP1 impair its regulatory ability, causing elevated expressions of NRF2 and NRF2-regulated gene transcription (24,27). It has been widely reported that increased expression of NRF2 causes resistance to therapies against malignant tumors (28–31).

Although *KEAP1* mutations are found in ∼15% of NSCLC patients, it is present in only about 2% of PDAC patients. However, *KEAP1* truncations were detected in tumors from PDAC patients who relapsed from daraxonrasib treatment (14). Moreover, KEAP1 inhibition is frequently found in malignant pancreatic cells through transcriptional or post-translational deregulation (31–33), and NRF2 overexpression has been observed as PDAC tumors progress to more aggressive stages (29,31). In a murine model of PDAC tumorigenesis, activation of Kras, Braf, or Myc induced Nrf2 expression, and genetic silencing of *Nrf2* impaired tumorigenic growth (34). In human PDAC cells, activated NRF2 promoted chemoresistance to gemcitabine (29). Collectively, these findings suggest that NRF2 plays an important role in PDAC tumor progression and regulating its sensitivity to treatments. However, the detailed mechanism regarding how the KEAP1-NRF2 pathway affects sensitivity to KRAS inhibition remains to be determined.

In this study, we explored the role of the KEAP1-NRF2 signaling pathway in mediating resistance to KRAS inhibition in KRAS-mutant PDAC and LUAD. We defined a *KEAP1*^KO^-dependent gene signature in PDAC and validated the association of this signature with reduced KRAS inhibitor sensitivity across PDAC and LUAD cancer cell lines, patient-derived xenograft (PDX) tumors, and patient clinical samples. Finally, we exploited the increased glutamine dependency caused by *KEAP1* loss and determined that concurrent pharmacologic inhibition of glutamine metabolism enhanced KRAS inhibitor activity in KRAS-mutant PDAC and LUAD.

## RESULTS

### Loss-of-function CRISPR Screen Identified the KEAP1-NRF2 Pathway in Promotion of Resistance to RAS Inhibition

To identify mechanisms of resistance to inhibitors of mutant KRAS, we focused our study on the most prevalent KRAS mutation, G12D. We first evaluated the anti-tumor activity of two inhibitors, the pan-RAS inhibitor RMC-7977 (analog of daraxonrasib) and KRAS^G12D^-selective inhibitor MRTX1133 (hereinafter referred to as RASi and G12Di, respectively) (9,19). We previously evaluated RASi specificity in a panel of matched mouse embryo fibroblasts (MEFs) with loss of all three *Ras* genes (RASless MEFs) (35), where proliferative capacity was restored with ectopic expression of human wild-type or mutationally activated *KRAS* or *BRAF* mutant genes (36). RASi exhibited comparable potent inhibition of KRAS effector signaling for all KRAS mutants evaluated as measured by inhibition of the phosphorylation (pERK) and activation of ERK (19), the key KRAS effector in PDAC. In contrast, G12Di exhibited potent inhibition of pERK in RASless MEFs expressing KRAS^G12D^ at 10 nM whereas pERK inhibition in KRAS WT cells was seen only at 500 nM (Supplementary Fig. S1A). An approximately 10-fold lower potency was seen against KRAS^G13D^, 50-fold lower activity against KRAS^Q61H^ and limited to no activity was detected against other KRAS mutants. No activity was seen in BRAF^V600E^ cells, supporting the KRAS-selectivity of ERK inhibition.

We then characterized the sensitivity of a panel of KRAS^G12D^-mutant PDAC cell lines (SW1990, Pa16C, Pa14C, and PANC-1) to RASi and G12Di. Both effectively reduced the levels of pERK as well as the activities of two ERK substrates, the phosphorylation of RSK (pRSK) and expression levels of total MYC, at comparably low nanomolar concentrations (Supplementary Fig. S1B and S1C).

To elucidate the mechanisms underlying resistance to KRAS inhibitors in PDAC cells, we carried out a loss-of-function CRISPR-Cas9 screen using a focused library targeting components of major oncogenic signaling networks (37) (Fig. 1A). We determined enrichment of sgRNAs targeting multiple genes encoding tumor suppressors, several of which were previously associated with acquired resistance to KRAS^G12C^ inhibition in patients (e.g., *PTEN, APC,* and *RB1*) (16,17,22,38) (Fig. 1B). *STK11* co-mutation was associated with early disease progression and shorter survival in NSCLC patients treated with G12C inhibitors (23,39). Conversely, we found depletion of sgRNAs targeting genes with gain-of-function mutations identified in patients who relapsed on KRAS^G12C^-selective inhibitor (G12Ci) treatment (e.g., *EGFR*, *ERBB2*, *BRAF*, *MYC, FGFR2* and *PIK3CA* (16,17,22,38). The identification of genes associated with relapsed patients support the validity of the screen to detect bona fide mechanisms of resistance.

**Figure 1.**
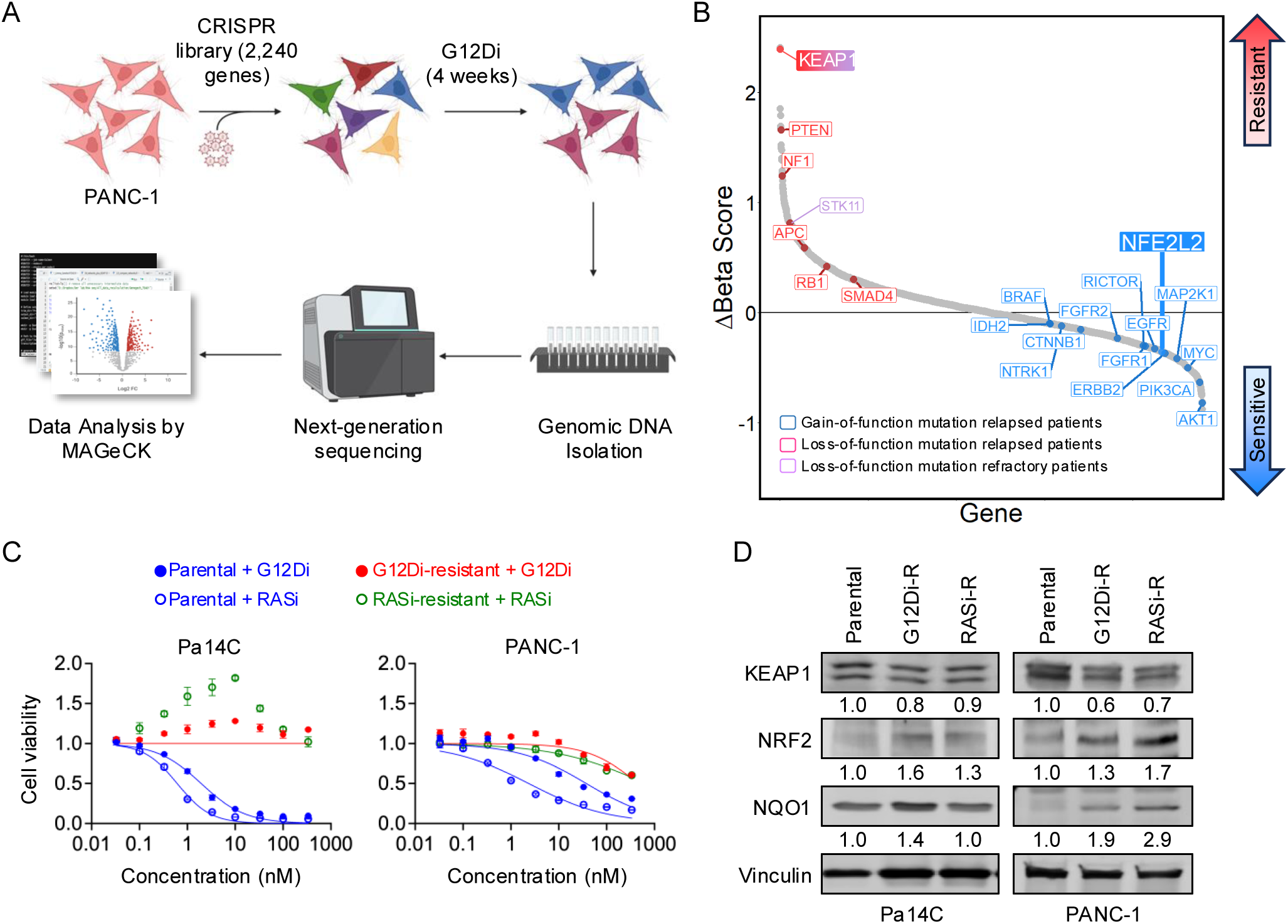
CRISPR screen on PANC-1 PDAC cell treated with G12Di identified the KEAP1-NRF2 pathway promoting resistance to KRAS inhibition. **A,** The loss-of-function CRISPR screen was performed using the druggable genome sgRNA library. The sgRNA library was transduced to cells by lentivirus and propagated growth medium supplemented with G12Di or vehicle (DMSO) control, for 4 weeks. The genomic DNA was isolated and sequenced by next generation sequencing (NGS). The NGS results were further analyzed by MAGeCK to determine the Beta scores for each gene. **B,** The snake plot highlights genes causing PANC-1 to be resistant or sensitized to G12Di identified in the CRISPR screens. Y axis represents the difference in Beta scores between G12Di- and DMSO-treated cells (Δ Beta Score). **C,** Resistant or parental PDAC cell lines were treated with G12Di (0 - 333 nM) or RASi (0 - 333 nM) for 5 days, and cell viability was determined by the addition of Hoechst. Normalized cell counts compared to the control conditions, which were normalized to 1.0, were plotted, and error bars denoted standard error of at least three independent biological replicates. **D,** Representative western blots probing KEAP1, NRF2, NQO1, and vinculin in Pa14C or PANC-1 cells which are resistant to G12Di (G12Di-R), RASi (RASi-R), or their parental counterpart. The relative levels of KEAP1, NRF2, and NQO1 expressions are denoted underneath the designated blot images.

Loss-of-function mutations in *KEAP1* were associated with shorter survival in NSCLC patients on G12Ci therapy (23,39), and we observed that *KEAP1* knockout was the most potent resistance driver. Conversely, *NFE2L2* (encoding NRF2/NFE2L2) knockout caused increased sensitivity to KRAS inhibitor treatment. These observations support a role for KEAP1 loss and NRF2 activation in causing resistance to KRAS inhibition.

To determine a role for KEAP1-NRF2 in acquired resistance, we cultured sensitive cell lines with each inhibitor to select for outgrowth of resistant subpopulations (Fig. 1C; Supplementary Fig. S1D). Whereas the GI_50_ values for each parental cell line for both inhibitors were ∼30 nM or less, the resistant cell populations exhibited very limited growth suppression at 100 nM or more of each inhibitor (Supplementary Fig. S1E). Western blot analysis determined that three of the four resistant cells exhibited lower KEAP1 and higher NRF2 expression levels, as well as increased expression of the NRF2 transcriptional target *NQO1* encoded protein (40) (Fig. 1D; Supplementary Fig. S1F). Together with the findings from our CRISPR screen, these results further support a role for the KEAP1-NRF2 signaling network in modulating sensitivity to KRAS inhibitors in KRAS-mutant PDAC.

### *KEAP1* Loss Induces NRF2-Dependent RAS Inhibitor Resistance in PDAC and LUAD

To validate the observations from the CRISPR screen, we applied CRISPR knockout of *KEAP1* in a panel of four KRAS^G12D^-mutant human PDAC cell lines (SW1990, Pa14C, Pa16C, and PANC-1) (Fig. 2A). An increase in the steady-state levels of NRF2 was seen in *KEAP1* knockout (*KEAP1*^KO^) relative to control cells stably infected with a lentivirus vector expressing Cas9 and sgRNA targeting eGFP. Increased expression of NRF2 transcription gene target encoded proteins (NQO1 and SLC7A11) was also seen, indicating activation of NRF2 transcriptional activity in the *KEAP1*^KO^ cell lines.

**Figure 2.**
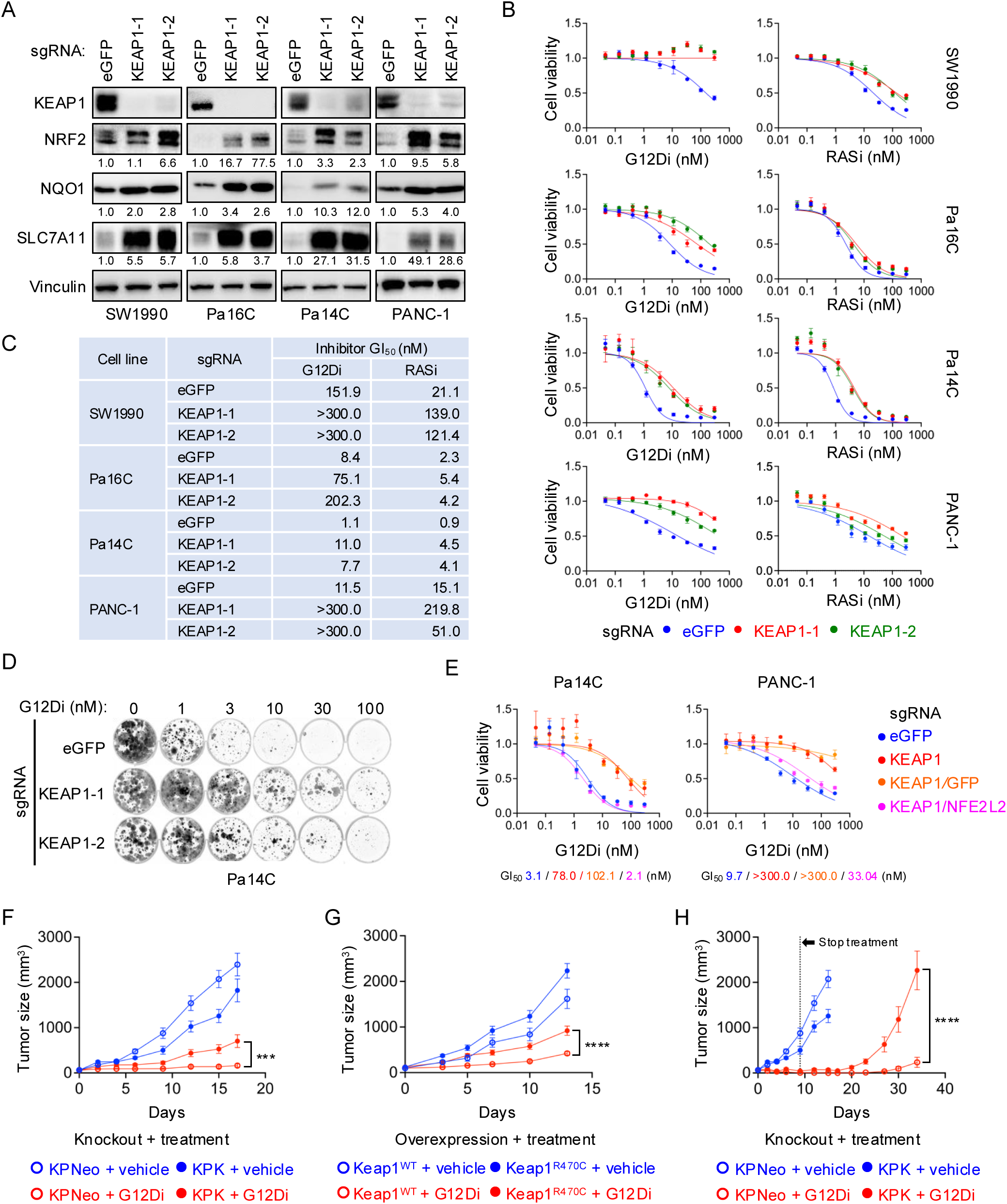
The knockout of KEAP1 drove resistance to KRAS inhibition by upregulating NRF2 expression. **A,** Representative western blots probing KEAP1, NRF2, NQO1, SLC7A11, and vinculin in PDAC cell lines (SW1990, Pa16C, Pa14C, and PANC-1) where *KEAP1* gene was knocked-out using two different sgRNA sequences targeting *KEAP1* (KEAP1-1 and KEAP1-2), or control sgRNA targeting *eGFP* (eGFP). The relative levels of NRF2, NQO1, and SLC7A11 expressions are denoted underneath the designated blot images. **B,** PDAC cell lines with *KEAP1* knockout (KEAP1-1, red; and KEAP1-2, green), or control cells (eGFP, blue), were treated with G12Di or RASi (0 – 300 nM) for 5 days, and cell viability was determined by the addition of Calcein-AM assay. Normalized cell counts compared to the control conditions, which were normalized to 1.0, were plotted, and error bars denoted standard error of at least three independent biological replicates. **C,** GI_50_ values for PDAC cells with *KEAP1* (KEAP1-1, KEAP1-2) or *eGFP* knockout treated with G12Di or RASi. **D,** Representative 2D clonogenic growth assays show long-term cell proliferation of Pa14C cells upon knockout of *KEAP1* (KEAP1-1 and KEAP1-2) or *eGFP*. Cells were cultured with G12Di (0 – 100 nM) for 14 days, and they were stained at the end of the experiment with crystal violet and imaged using a Typhoon FLA9500 biomolecular imager. **E,** PDAC cell lines (Pa14C and PANC-1) cells with knockout of *KEAP1* using KEAP1-1 sgRNA (KEAP1, red), double knockout of *KEAP1* and *GFP* (KEAP1/GFP, orange), double knockout of *KEAP1* and *NRF2* (KEAP1/NFE2L2, magenta), and control cells with *eGFP* knockout (eGFP, blue) were treated with G12Di (0 – 300 nM) for 5 days, and cell viability was determined by the addition of Calcein-AM assay. Normalized cell counts compared to the control conditions, which were normalized to 1.0, were plotted, and error bars denoted standard error of at least three independent biological replicates. Concentrations of G12Di inhibit 50% of viability were defined as GI_50_ and are shown in the corresponding colors underneath each plot. **F,** Tumor size measurements of mice bearing KP allografts with knockout of *Keap1* (KPK, solid dots) or neomycin resistant gene (KPNeo, hollow dots). Mice were treated with either vehicle (blue) or G12Di (10 mg/kg, red) and monitored for 17 days. **G,** Tumor size measurements of mice bearing KP allografts ectopically expressing wild type *Keap1* (Keap1^WT^, hollow dots) or *KEAP1*^R470C^ (KEAP1^R470C^, solid dots). Mice were treated with either vehicle (blue) or G12Di (10 mg/kg, red) and monitored for 13 days. **H,** Tumor size measurements of mice bearing KP allografts with knockout of *Keap1* (KPK, solid dots) or neomycin resistant gene (KPNeo, hollow dots). Mice were treated with either vehicle (blue) or G12Di (30 mg/kg, red) for 9 days. The tumor growth was monitored for 34 days or until they reach the sacrificing endpoints.

We found that *KEAP1*^KO^ cells exhibited variable resistance to both G12Di and RASi with significantly higher GI_50_ values relative to *eGFP*^KO^ control cells (Fig. 2B and 2C). *KEAP1*^KO^ cells also exhibited several fold reductions (1.8- to 2.6-fold) in G12Di induced cell death (Supplementary Fig. S2A). *KEAP1*^KO^ cells exhibited reduced sensitivity to G12Di treatment, compared to *eGFP*^KO^ control cells in long-term clonogenic growth assays (Fig. 2D; Supplementary Fig. S2B and S2C). Similarly, *KEAP1*^KO^ cells showed reduced sensitivity to G12Di treatment when cultured in three-dimensional (3D) non-adherent spheroid cultures (Supplementary Fig. S2D).

To determine whether the resistance induced by the loss of *KEAP1* is dependent on NRF2 expression, we generated *KEAP1* and *NFE2L2* double knockout (*KEAP1/NFE2L2*^KO^) PANC-1 and Pa14C cells together with control *KEAP1*/*GFP*^KO^ knockout cell lines. Compared with *KEAP1*/*GFP*^KO^ cells, the *KEAP1/NFE2L2*^KO^ cells showed decreased expression of NQO1, verifying reduction in NRF2-mediated transcriptional activity (Supplementary Fig. S2E). The *KEAP1*/*GFP*^KO^ control cells exhibited similar resistance to G12Di as *KEAP1*^KO^ cells, whereas *KEAP1/NFE2L2*^KO^ cells showed comparable sensitivity as control PDAC cells (Fig. 2E). We concluded that resistance to RAS inhibition driven by *KEAP1* loss was dependent on NRF2 expression.

We also assessed the consequences of pharmacologic induction of NRF2 expression using two chemically distinct small molecules, AI-1 and bardoxolone methyl (CDDO-me), which block KEAP1 interaction with NRF2, stabilizing NRF2 expression (41,42). Treatment with either AI-1 or CDDO-me caused increased expression of NRF2 (Supplementary Fig. 2F) and reduced sensitivity to G12Di (Supplementary Fig. 2G). Taken together, these observations suggest that NRF2 activation is both necessary and sufficient to support resistance to RAS inhibition.

We then extended our analyses of KEAP1 loss in driving resistance to RAS inhibition in LUAD. We first knocked out *KEAP1* in the KRAS^G12V^-mutant LUAD cell line NCI-H441 by CRISPR-Cas9 editing. Similar to what we observed in PDAC cell lines, *KEAP1*^KO^ resulted in upregulation of NRF2 expression and transcriptional activity, and resistance to RASi (Supplementary Fig. S2H and S2I). Second, using murine *Kras*^G12D^/*Trp53*^−/−^ (KP) cells with *Keap1* knockout (KPK) (43), we established KPK cells ectopically expressing wild-type *Keap1* or an empty vector control. Restoration of Keap1 expression caused a five-fold increased sensitivity to G12Di (Supplementary Fig. S2J). Together, these observations support KEAP1 loss as a mechanism of resistance to RAS inhibitors in LUAD.

We then investigated the impact of *Keap1* loss on response to G12Di in syngeneic immune competent tumor models established by KPK or KP cells with a control CRISPR-Cas9 knockout (KPNeo) (43,44). Tumor growth was monitored in mice treated with G12Di or vehicle. We found that *Keap1* loss significantly reduced tumor response to G12Di (Fig. 2F). Similarly, we treated KP cells with ectopic expression of either wild type (KPK^WT^) or a dominant negative *Keap1*^R470C^ mutant with reduced ability to degrade NRF2 (KPK^R470C^) with G12Di and observed that tumors expressing Keap1^R470C^ grew significantly faster than those expressing Keap1^WT^ when treated with G12Di (Fig. 2G). Finally, we tested the durability of G12Di response by treating mice bearing KPK or KPNeo allografts with G12Di or vehicle for 9 days, after which we stopped the treatment (Fig. 2H). While G12Di caused robust tumor suppression in both models, KPK tumors rebounded after day 20 and grew rapidly. However, the KP control tumors remained suppressed and showed limited signs of relapse until day 30. We conclude that loss of *KEAP1* can drive resistance to RAS inhibition in both PDAC and LUAD *in vitro* and *in vivo*.

### KEAP1 Loss Drives a Distinct Transcriptome Independent of Oncogenic KRAS

Patients who relapsed on G12Ci therapy exhibit mutational activation of ERK signaling (16,17,22,38), and we showed that ERK reactivation alone was sufficient to drive RAS inhibitor resistance (19,45,46). Therefore, we examined if *KEAP1* loss-driven G12Di resistance involved activation of ERK. Control (*eGFP*^KO^) and *KEAP1*^KO^ matched pairs of PANC-1 and Pa14C cell lines were treated with G12Di, and the suppression of ERK signaling was monitored (Supplementary Fig. S3A). Comparable suppression of pERK and the ERK substrate RSK (pRSK) was seen after 4- and 24-hour G12Di treatment regardless of the *KEAP1* status. We conclude that *KEAP1* loss-induced resistance was not due to an impaired ability of G12Di to inhibit ERK.

To assess a molecular basis for *KEAP1* loss-mediated resistance against G12Di, we conducted RNA-sequencing (RNA-seq) to profile gene expression changes in four PDAC cells depleted of *KEAP1*. We applied a doxycycline (dox)-inducible CRISPR-Cas9 system (47) to acutely (5 days) knockout *KEAP1* in four *KRAS*^G12D^-mutant PDAC cell lines. *KEAP1* knockout caused elevated expression of NRF2 and NRF2 transcriptional targets *NQO1* and *SLC7A11* (Supplementary Fig. 3B) and promoted resistance to G12Di to similar levels seen with stable *KEAP1* knockout (Supplementary Fig. S3C).

Our RNA-seq analyses detected a total of 17,674 genes (with read counts > 20) across all four cell lines. Among these, 1,248 genes were found significantly upregulated (PDAC-*KEAP1*^KO^ UP, log_2_ fold change (FC) > 0, adjusted *p* value (adj. *p*) < 0.05) and 1,016 genes significantly downregulated (PDAC-*KEAP1*^KO^ DN, log_2_FC < 0, adj. *p* < 0.05) upon *KEAP1* knockout (Fig. 3A; Supplementary Table S1).

**Figure 3.**
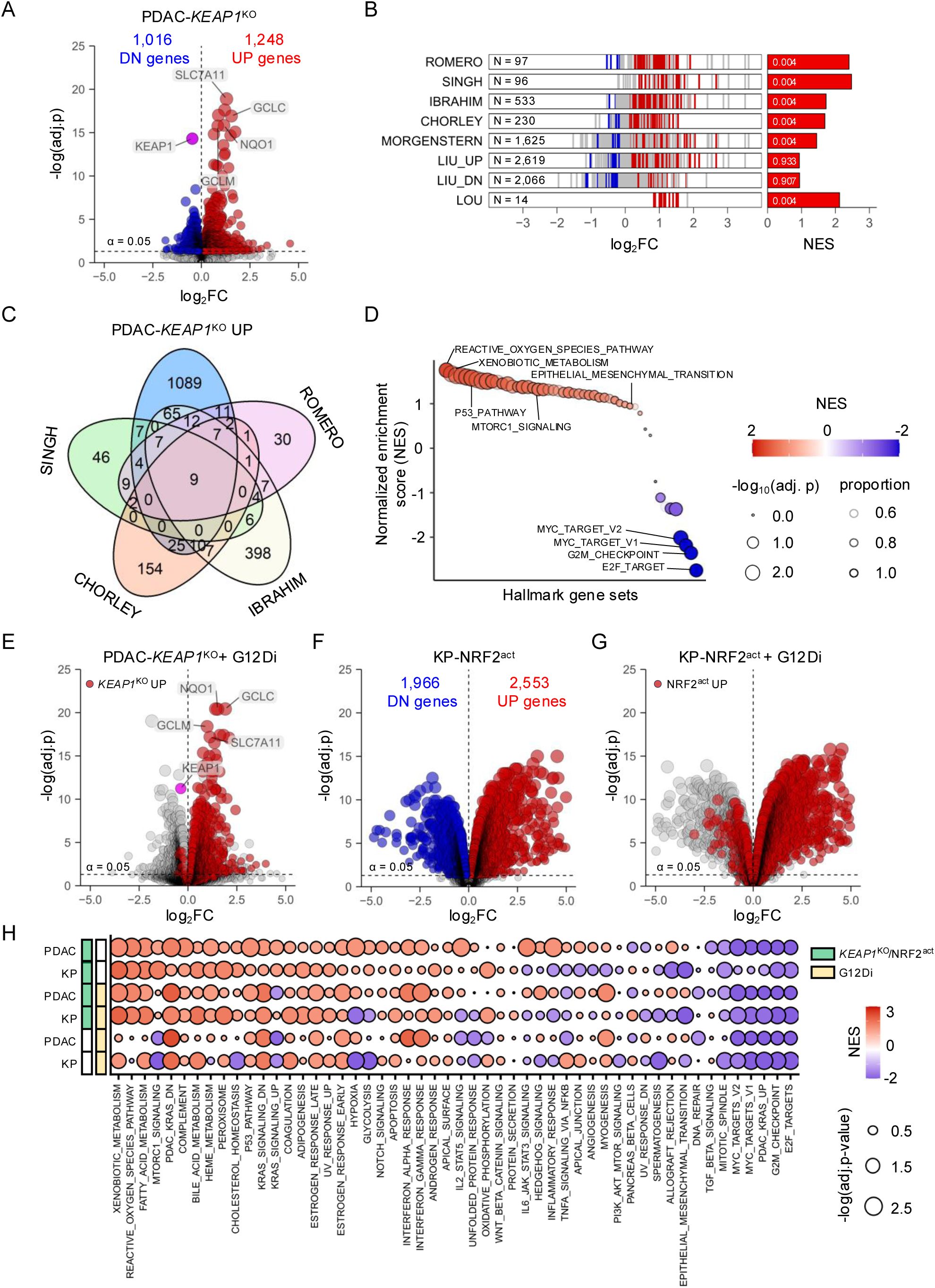
RNA sequencing identified unique KEAP1-loss transcriptome independent of KRAS. **A,** Volcano plots showing the differentially expressed mRNA transcripts from the RNA-seq analysis in PDAC cells following induced *KEAP1* knockout, the PDAC-*KEAP1*^KO^ transcriptome. A total of 1,248 significantly up-regulated (red) and 1,016 significantly down- regulated (blue) genes were identified. Representative NRF2 transcriptional targets, including SLC7A11, GCLC, NQO1, and GCLM were found significantly upregulated in the KEAP1-loss transcriptome. **B,** Barcode plots illustrating the differential expression (log_2_FC) in the PDAC-*KEAP1*^KO^ transcriptome of genes within each signature (left panel). Significantly upregulated and downregulated genes within each signature are shown in red and blue, respectively. The number of genes (N) in each set is labeled in each box (right panel) Normalized enrichment scores (NES) for the corresponding signatures of the PDAC-*KEAP1*^KO^ transcriptome in (A). Adjusted P values are indicated within the bars. **C,** Venn diagram showing the numbers of overlapping genes within ROMERO, IBRAHIM, CHORLEY, SINGH signatures, and the PDAC-*KEAP1*^KO^ UP signature from (A). **D,** Dot plot showing normalized enrichment scores (NES) of 50 Hallmark gene sets from GSEA on the PDAC *KEAP1*^KO^ transcriptome. **E,** Volcano plots showing the differentially expressed mRNA transcripts from RNA-seq analysis on *KEAP1*^KO^ PDAC cells treated with G12Di at their GI_50_ concentrations for three days. 65% of PDAC-KEAP1^KO^ UP genes remain significantly upregulated (red). **F,** Volcano plots showing the differentially expressed mRNA transcripts from RNA-seq analysis on KP mouse lung cancer cells treated with NRF2 activator KI696 (500 nM) for 6 days. A total of 2,533 significantly up-regulated (red) and 1,966 significantly down-regulated (blue) genes were identified. **G,** Volcano plots showing the differentially expressed mRNA transcripts from RNA-seq analysis of KP cells pre-treated with NRF2 activator KI696 (500 nM) for 4 days and G12Di (10 nM) for 2 days. Genes upregulated in (F) remain significantly upregulated under KRAS inhibition (red). **H,** GSEA performed using the 50 Hallmark gene sets, as well as PDAC-KRAS UP and DN signatures previously reported by our group. Analyses were conducted on PDAC cell panels (PDAC) and KP LUAD cells (mKP) with or without *KEAP1* knockout (*KEAP1*^KO^) or NRF2 activation (NRF2^act^), and with or without G12Di treatment.

We first performed HOMER (Hypergeometric Optimization of Motif EnRichment) analyses to assess the transcription factor binding motifs present in the PDAC-*KEAP1*^KO^ UP and DN gene sets (48). Supervised promoter motif-enrichment analyses for the top 500 UP gene set identified the DNA motif for NFE2L2/NRF2 as the top motif (Supplementary Fig. S3D), and this motif corresponds to the antioxidant response element (ARE) (49). In contrast, no ARE motifs were identified in the top 10 scoring motifs for the PDAC-*KEAP1*^KO^ 500 DN genes (Supplementary Fig. S3E). Given that we determined that *KEAP1* loss-associated resistance to RASi was NRF2-dependent (Fig. 2E), we speculated that the PDAC-*KEAP1*^KO^ UP but not DN gene set comprises the significant genes involved in driving RASi resistance.

We next compared our PDAC-*KEAP1*^KO^ UP/DN 2,264-gene signature with eight published gene signatures previously reported by KEAP1 loss and/or NRF2 activation in rodent and/or human species (Supplementary Table S2) (43,50–55). Although gene set enrichment analysis (GSEA) found enrichments in six of these KEAP1-NRF2 signatures in our PDAC-*KEAP1*^KO^ RNA-seq, only our previously established ROMERO signature (43) and the 14-gene LOU signature showed substantial overlaps with our gene set (49 and 93%, respectively). In contrast, fewer than 30% of genes within the remaining five signatures were significantly upregulated in our gene set (Supplementary Fig. S3F; Fig. 3B).

Reflecting the distinct nature of our PDAC-*KEAP1*^KO^ UP gene signature, we found only nine genes overlapping with the ROMERO, SINGH, IBRAHIM, and CHORLEY signatures (Fig. 3C). In contrast, 1,089 genes in our PDAC-*KEAP1*^KO^ UP gene set (87%) have not been identified previously in any of these published gene sets. The lack of overlap with previous signatures may reflect the distinct experimental approaches for activating NRF2 as well as the different cell models studied, with none *KRAS*-mutant or pancreatic cell models.

We performed overrepresentation analysis (ORA) using the mSigDB 50 Hallmark gene sets and found that the p53, hypoxia, mTORC1 signaling, reactive oxygen species, and glycolysis pathway gene sets were highly enriched in our PDAC-*KEAP1*^KO^ UP signature as well as in the published KEAP1-NRF2 gene sets. The convergence of these signaling pathways underscores a conserved core of NRF2-mediated functions, highlighting the robustness of our signature in capturing KEAP1-NRF2 oncogenic function (Supplementary Fig. 3G).

We next evaluated the 1,089 unique genes within the PDAC-*KEAP1*^KO^ UP gene set by ORA using Reactome, Kyoto Encyclopedia of Genes and Genomes (KEGG), and Gene Ontology (GO) gene sets. There were nine pathways that were found significantly enriched, with five of these related to lipid metabolism or biosynthesis (Supplementary Fig. 3H). Moreover, HOMER analysis performed on these 1,089 unique genes no longer showed any enriched transcription binding motifs, including NFE2L2/NRF2 (Supplementary Fig 3I). These results suggest that secondary gene targets regulated by the KEAP1-NRF2 pathway were captured in our transcriptomic analysis. These analyses highlight that our PDAC-*KEAP1*^KO^ model captures KEAP1-NRF2-associated genes distinct from the core NRF2 gene targets.

To assess the biological processes regulated by KEAP1 in PDAC, we evaluated our PDAC KEAP1-dependent transcriptome by gene set enrichment analyses (GSEA) with the 50 Hallmark gene sets. As expected, the REACTIVE_OXYGEN_SPECIES_PATHWAY was among the top upregulated activity (Fig. 3D).

To evaluate whether the RAS inhibition can suppress *KEAP1*^KO^ transcriptome, we performed RNA-seq on these PDAC cells cultured with doxycycline to induce KEAP1 knockout and treated with G12Di. Interestingly, only 11 genes (< 1%) within the 1,248-gene PDAC-*KEAP1*^KO^ UP signature were suppressed by G12Di, while 813 genes (65%) of the signature remained upregulated under G12Di treatment (Fig. 3E). Moreover, we extended our RNA-seq analysis again using KP mouse lung cancer cells cultured in 3D spheroid and treated with a NRF2 inducer KI696 (KP-NRF2^act^) to induce NRF2 expression (56). We detected a total of 13,182 genes (11,079 genes mapped with human homologs), with 2,553 upregulated and 1,966 downregulated genes (Fig. 3F). Similarly, 76 % (1,931) gene expressions that were upregulated upon KI696 treatment sustained upon treated with G12Di, with only 9% (232) gene exhibited decreased expressions (Fig. 3G). Overall, these observations suggest that KEAP1-NRF2 regulated signaling pathways that are independent of KRAS-regulated signaling networks.

We next performed GSEA with the MSigDB 50 Hallmark gene sets plus our previously established PDAC-KRAS UP/DN gene signature on PDAC-*KEAP1*^KO^ or KP-NRF2^act^ treated without or with G12Di, and control PDAC or KP cells treated with G12Di as controls (45,57). Trends for gene sets showing either upregulation or downregulation were generally consistent across all PDAC and KP models (Fig. 3H, and Supplementary Fig. S3J). As expected, the most significant activities upregulated in *KEAP1*^KO^ cells were the Reactive Oxygen Species Pathway and the Xenobiotic Metabolism gene sets, well-established NRF2-driven activities (24,27), and these pathways remain enriched even when cells were treated with G12Di. Importantly, although G12Di cause a significant downregulation of the PDAC-KRAS UP gene set, *KEAP1* loss or NRF2 activation cannot restore its expression, consistent with our observation that *KEAP1*^KO^ was not able to restore the activation of ERK from G12Di (Supplementary Fig. S3A). Furthermore, we performed ORA on genes that are significantly upregulated by more than 50% (adj. *p* < 0.05, FC > 1.5) for each cell line (Supplementary Fig. 3K), and we determined that 29 gene sets that were significantly upregulated in at least three of four cell lines. These gene sets included the Reactome KEAP1-NFE2L2 pathway, Nuclear events mediated by the NFE2L2 pathway, and NFE2L2 regulating antioxidant/detoxification enzymes gene sets, which is consistent with our findings that *KEAP1* depletion activated NRF2-dependent gene transcription.

The sustained activation of the Reactive Oxygen Species Pathway in *KEAP1*^KO^ or NRF2^act^ cells treated with G12Di (Fig. 3H) identified one mechanistic basis for resistance to RAS inhibition. A previous study found that treatment of *BRAF*-mutant lung or *KRAS*-mutant pancreatic cancer cell lines with the MEK inhibitor trametinib (MEKi) induced accumulation of ROS, but this effect was attenuated with *KEAP1* loss (30). Therefore, we determined whether KRAS-ERK inhibition causes oxidative stress by accumulating ROS, and whether *KEAP1* knockout would counter ROS activation. We treated control (*eGFP*^KO^) or *KEAP1*^KO^ PANC-1 and Pa14C cells with G12Di or MEKi, and the level of ROS was determined using the CellROX Green fluorogenic probe. Both G12Di and MEKi treatment increased ROS levels in control cells but not *KEAP1*^KO^ cells, except for *KEAP1*^KO^ PANC-1 cells which exhibited a limited ROS increase when treated with MEKi (Supplementary Fig. S3L).

It has been observed that KRAS inhibitors and oxidative stress can induce cell cycle arrest, and previous studies established that NRF2 promotes cell cycle transition (58–61). Therefore, we examined whether *KEAP1* knockout and NRF2 accumulation could mitigate cell cycle arrest induced by KRAS inhibition. We performed cell cycle analyses on *KEAP1*^KO^ and control (*eGFP*^KO^) PANC-1 and Pa14C cells treated with G12Di for three days. Control Pa14C cells showed a pronounced G_1_ arrest upon treatment with G12Di, while *KEAP1*^KO^ Pa14C cells only exhibited modest increase in G1 population under G12Di treatment (Supplementary Fig. S3M). Similarly, in PANC-1, G12Di caused an increase in G_1_ population, while negligible changes in cell cycle distribution were observed in *KEAP1*^KO^ PANC-1. These findings suggest that a subset of *KEAP1*^KO^ PDAC cells are refractory to cell cycle arrest induced by KRAS inhibition.

### Distinct Mechanisms of Resistance Due to Loss of KEAP1 Versus Activation of KRAS-MYC or YAP/TAZ-TEAD

MEK-ERK, MYC, and YAP/TAZ-TEAD activation has been reported to limit the effectiveness of KRAS inhibition (12,17,20,22). To assess the degree of overlap between genes regulated by these mechanisms of resistance with KEAP1-NRF2, we compared the *KEAP1*^KO^ UP transcriptome to transcriptomes associated with KRAS, MYC and YAP/TAZ-TEAD. For these analyses, we utilized data from gene sets established by acute 24 hour siRNA knockdown of *KRAS* (KRAS UP) (45) or *MYC* (MYC UP) (62) in a panel of eight *KRAS*-mutant PDAC cell lines, or pharmacologic inhibition with 48 hour treatment of an *NF2*-null human mesothelioma cell line with the pan-TEAD inhibitor GNE-7883 (TEAD UP; GSE273175) (63). We performed the network analysis on genes that were significantly upregulated from each dataset and weighted based on their weighted log_2_FC (Supplementary Fig. S4A).

We found very significant overlap with the KRAS-, MYC-, and TEAD-dependent gene networks (607 genes), while the *KEAP1*^KO^-dependent transcriptome was distinct from the other three gene sets, with only 4-13% overlap (Fig. 4A). Moreover, analysis of gene essentiality scores in pancreatic and lung cancers from the Dependency Map (DepMap) portal shows that the KRAS-, MYC-, and TEAD-dependent genes generally exhibited higher essentiality scores, whereas the *KEAP1*^KO^-dependent UP transcriptome was comprised primarily of non-essential genes. Collectively, these findings suggest that the KEAP1-NRF2 signaling network promotes drug resistance utilizing a distinct molecular mechanism from those driven by KRAS, MYC, or TEAD.

**Figure 4.**
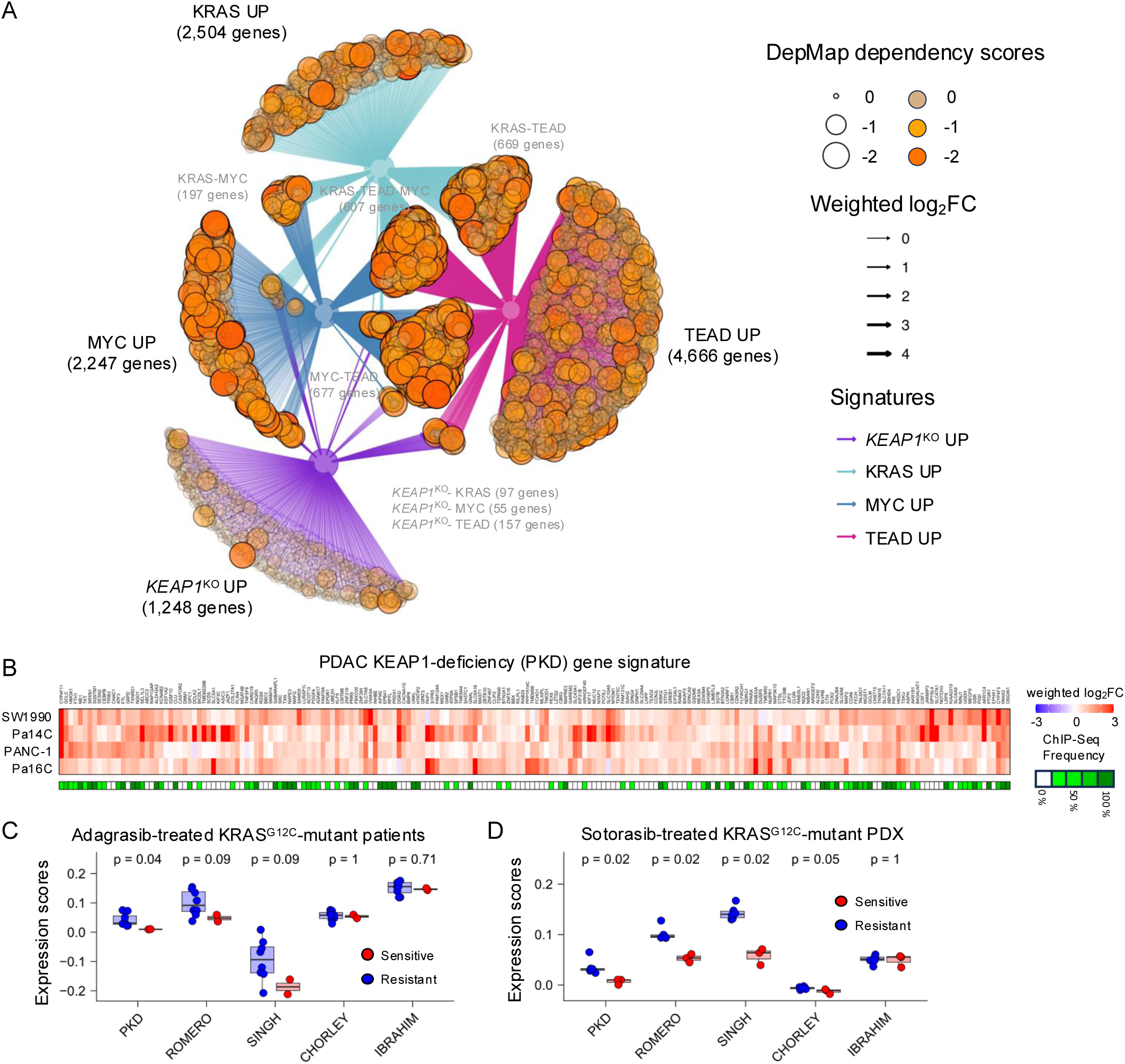
KEAP1-loss transcriptome is distinct from KRAS-, TEAD-, and MYC-dependent transcriptome and associated with the resistance to KRAS inhibitors. **A,** The network analysis on genes upregulated upon KEAP1 knockout (*KEAP1*^KO^ UP), as well as genes dependent on KRAS (KRAS UP), MYC transcription factor (MYC UP), or TEAD transcription factor (TEAD UP). Each node denotes a gene, and the size and color intensity for each node represent the medium DepMap essentiality scores across PDAC and LUAD cohorts. The edges are color coded and connect each node to their associated regulatory programs (KRAS, TEAD, MYC, or *KEAP1*^KO^). The width of each edge represents the magnitude of weighted log_2_FC from each dataset. **B,** The heatmap for the expression changes (weighted log_2_FC) of genes within the PKD signature. The frequency (ChIP-Seq frequency, green) of genes that were found to be NRF2 transcriptional targets in the ENCODE database were plotted underneath the plot. **C,** The gene expression scores of PKD, ROMERO, SINGH, CHORLEY, and IBRAHIM signatures were determined using Singscore on mRNA expressions collected from KRAS^G12C^-mutant patients before treated with adagrasib. The p values between patients who responded (sensitive, red) or were resistant to adagrasib (resistant, blue) were determined using Wilcoxon test. **D,** The gene expression scores of PKD, ROMERO, SINGH, CHORLEY, and IBRAHIM signatures were determined using Singscore on mRNA expressions collected from PDX models treated with sotorasib. The p values between initially sensitive tumors (sensitive, red) or relapsed tumors (resistant, blue) were determined using Wilcoxon test.

### Establishment of a PDAC KEAP1-Deficiency (PKD) Gene Signature

With *KEAP1* loss associated with RASi resistance and deregulated gene transcription, we next determined a KEAP1-deficiency gene signature that may have predictive value for response to RASi. To define a focused gene set, we utilized the top 200 significantly upregulated and protein-coding genes upon *KEAP1*^KO^ as the PDAC KEAP1-deficiency (PKD) gene signature (Fig. 4B; Supplementary Table 3). We first examined the proportion of direct NRF2 transcriptional targets within the PKD signature by cross-referencing chromatin immunoprecipitation sequencing (ChIP-seq) data from the ENCODE database (64). We found 100 genes (50%) within the PKD signature were detected as direct NRF2 gene targets (Fig. 4B). Moreover, HOMER motif analysis identified NFE2L2, BACH1, and NRF2 as top enriched transcription factor binding motifs in the PKD signature, where BACH1 has been reported to act as a repressor to AREs as a negative feedback mechanism of NRF2 activation (Supplementary Fig. S4B) (65). In summary, the PKD signature captures a significant portion of direct NRF2 transcriptional targets as well as secondary targets where their genetic expressions were induced upon *KEAP1*^KO^.

To elucidate the pathways regulated by genes within the PKD signature, ORA was carried out to identify significantly enriched pathways using Reactome, KEGG, and GO gene sets (Supplementary Fig. S4C). The analysis revealed a robust enrichment associated with oxidative stress response pathways. Given our previous finding that KRAS inhibition elevates ROS levels in PDAC cells (Supplementary Fig. S3L), these ORA results underscore the important role of oxidative stress relief of the PKD genes.

Beyond oxidative stress control, the PKD signature consists of genes potentially involved in additional mechanisms promoting resistance to KRAS inhibitors. ORA revealed that pathways related to chemical stress or toxic substance response were highly enriched. These include *ABCC3* and *SLC47A1*, which encode membrane transporters that facilitate drug efflux, diminishing inhibitor efficacy (66–68). Aldo-keto reductase genes, such as *AKR1B1* and *AKR1C4*, were identified, and they have been shown to confer chemoresistance and resistance to small molecule inhibitors via NADPH-dependent carbonyl reduction to external chemicals (69–72). Autophagy related genes (*TXNRD1* and *SQSTM1*) were further notable components, and we and others previously reported that increased autophagy led to resistance to ERK inhibition in PDAC cells (73,74). The *SRC* tyrosine kinase oncogene was also identified in the PKD signature and is frequently overexpressed in PDAC and has been widely reported to cause drug resistance (75–77).

Finally, we examined the clinical relevance of the PKD signature in LUAD patients harboring mutant KEAP1. Utilizing RNA-seq data from the LUAD cohort in TCGA database, we categorized patients based on their *KEAP1* mutational status and calculated the enrichment scores of genes within the PKD signature. The *KEAP1* mutant tumors exhibited significantly elevated expressions of the PKD signature relative to those carrying wild type *KEAP1* (Supplementary Fig. S4D). These data further confirm that the PKD gene set is a robust indicator of aberrant KEAP1 signaling in not only PDAC but also LUAD.

### Increased Expression of Genes within the PKD Signature is Associated with Resistance to KRAS Inhibition

We next determined if the PKD signature is associated with resistance to KRAS inhibitors. We examined gene expressions within the PKD, ROMERO, SINGH, CHORLEY, and IBRAHIM signatures in the RNA expression data from patients or patient-derived xenografts (PDX) treated with KRAS inhibitors. The transcriptome of KRAS^G12C^-mutant NSCLC or CRC patients before treatment with adagrasib (45) showed that the two patients who responded to the drug had the lowest PKD expression scores, and those were significantly separated from the non-responder group (Fig. 4C). In contrast, expression scores of ROMERO, SINGH, CHORLEY, and IBRAHIM signatures did not effectively differentiate the response status.

A similar analysis was performed on PDX models treated with sotorasib (GSE204753) (78), and we found the significantly lower expression scores of PKD, ROMERO, and SINGH signatures in the sensitive samples, distinguishing them from those which were resistant (Fig. 4D). The CHORLEY signature showed limited distinction while IBRAHIM expression scores were indistinguishable between the two groups.

In summary, analyses across patient cohorts and PDX models under RASi treatments demonstrated that the PKD gene signature was strongly associated with resistance to KRAS inhibitors and poor clinical outcomes. We conclude that, although other previously established gene sets encompass a broader coverage of KEAP1-NRF2-related genes, the PKD signature can more effectively distinguish RASi resistant cases from sensitive samples.

### *KEAP1*^KO^ PDAC cells Exhibit Enhanced Dependency on Glutamine Metabolism

We have previously demonstrated that *KEAP1*^KO^ lung cancer cells become more sensitive to the glutaminase inhibitor CB-839 (GLSi) (43). Glutaminase facilitates the conversion of glutamine to glutamate, and it is critical for glutathione synthesis and metabolic reprogramming in cancer cells (79,80). In fact, we and others have reported that cells with KEAP1 inactivation or NRF2 activation can reduce oxidative stress by increasing glutathione levels (26,30,43,81). In the present study, we found that the expressions of genes regulating glutamine and glutathione metabolism were enriched in the cancer cells with *KEAP1* loss, such as *PGD*, *G6PD*, *GSR*, *GCLC*, and *GCLM*. This finding led us to investigate whether *KEAP1* loss in PDAC cells is associated with increased dependency on glutamine metabolism.

We evaluated cell proliferation upon treating control and *KEAP1*^KO^ PDAC cells with GLSi. We first applied a glutaminase activity assay using isolated mitochondria and confirmed the inhibitory effect of GLSi on glutaminase activity (Supplementary Fig. S5A). The proliferation assay performed on SW1990, Pa16C, and Pa14C PDAC showed enhanced sensitivity to glutaminase inhibition upon KEAP1 knockout, while the sensitivity of GLSi changed only marginally in PANC-1 cells (Fig. 5A). We then examined glutamate secretion and glutamine consumption in *KEAP1*^KO^ SW1990 and Pa14C cell lines. We found that glutamate secretion and glutamine uptake significantly increased in the *KEAP1*^KO^ cells (Supplementary Fig. S5B and S5C), confirming elevated glutamine dependency.

**Figure 5.**
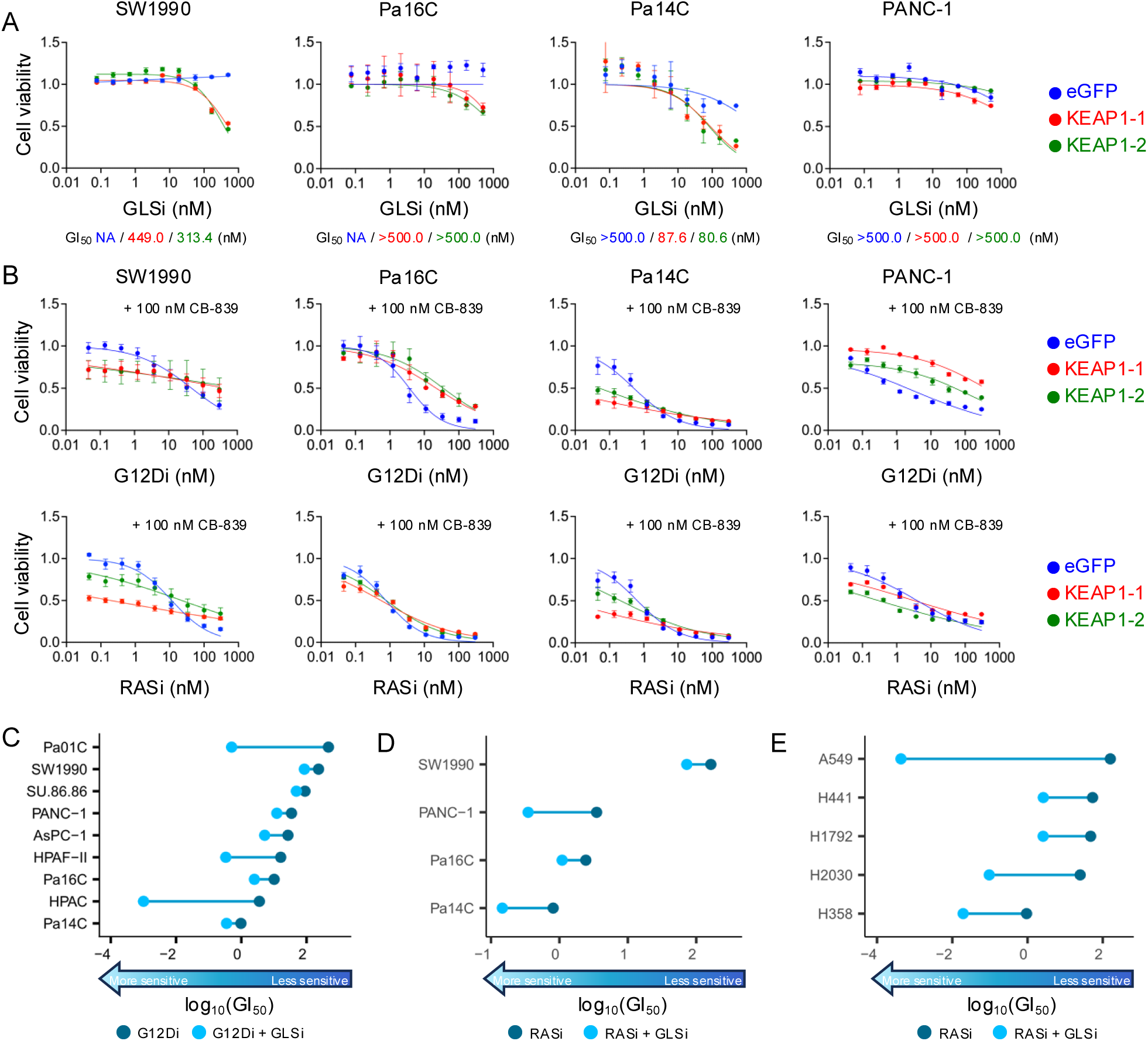
Concurrent inhibition of glutaminase and KRAS showed enhanced anti-tumor activity in PDAC. **A,** PDAC cell lines with stable knockout of *KEAP1* (KEAP1-1, red; and KEAP1-2, green), or *eGFP* (eGFP, blue), were treated with GLSi (0 - 500 nM) for 5 days, and cell viability was determined by the addition of Calcein-AM assay. Normalized cell counts compared to the control conditions, which were normalized to 1.0, were plotted, and error bars denoted standard error of at least three independent biological replicates. Concentrations of GLSi that inhibit 50% viability were defined as GI_50_ and are shown in the corresponding colors underneath each plot. **B,** PDAC cell lines with stable knockout of *KEAP1* (KEAP1-1, red; and KEAP1-2, green), or *eGFP* (eGFP, blue), were treated with GLSi (100 nM) and either G12Di (top panel, 0 - 300 nM) or RASi (bottom panel, 0 - 300 nM) for 5 days, and cell viability was determined by the addition of Calcein-AM assay. Normalized cell counts compared to the control conditions, which were normalized to 1.0, were plotted, and error bars denoted standard error of at least three independent biological replicates. **C,** The GI_50_ (nM) of PDAC cell panel treated with G12Di, and without (dark blue) or with (light blue) 100 nM GLSi. **D,** The GI_50_ (nM) of PDAC cell panel treated with RASi. and without (dark blue) or with (light blue) 100 nM GLSi. **E,** The GI_50_ (nM) of LUAD cell panel treated with RASi, and without (dark blue) or with (light blue) 100 nM GLSi.

Next, we investigated the mechanism underlying the enhanced glutamine dependency caused by *KEAP1* depletion. Glutamate secretion is mediated by the anti-transporter SLC7A11, which exchanges intracellular glutamate for extracellular cystine, another precursor for glutathione synthesis (43,82,83). We showed previously that SLC7A11 overexpression is associated with increased sensitivity to glutaminase inhibition (43), and in the present study we determined that *SLC7A11* mRNA and protein expression were upregulated upon *KEAP1* depletion (Fig. 2A and 3A). To determine whether *KEAP1*^KO^-associated glutamine addiction resulted from SLC7A11 overexpression in PDAC cells, we knocked down *SLC7A11* with two distinct siRNAs (siSLC7A11-1 and siSLC7A11-2), and with non-specific siRNA as controls (siNS), in control (*eGFP*^KO^) or *KEAP1*^KO^ SW1990 and PA14C cells. The reduction of SLC7A11 expression (Supplementary Fig. S5D) resulted in decreased sensitivity to GLSi in *KEAP*^KO^ cells (Supplementary Fig. S5E) (43). Collectively, these findings support a mechanism where *KEAP1*-depleted PDAC cells become dependent on glutaminase activity primarily through upregulation of SLC7A11 expression.

We next determined if combining GLSi with a KRAS inhibitor can more effectively inhibit cancer cell proliferation. With the addition of GLSi, we found that the GI_50_ of G12Di or RASi became notably lower in *KEAP1*^KO^ cells, and this combination also enhanced the cytotoxic activity of KRAS inhibitors in control cells (Fig. 5B; Supplementary Fig. S5F and S5G). We also evaluated the sensitivity of G12Di or RASi without or with GLSi in a panel of primary PDAC and NSCLC cell lines. We found that concurrent treatment with GLSi generally augmented the activity of G12Di and RASi across PDAC and LUAD cell lines, with Pa01C, HPAF-II, HPAC, and A549 cells presented the strongest enhancement (Fig. 5C-5E; Supplementary Fig. S5H-L). These observations support the benefit of co-targeting glutamine metabolism and KRAS signaling in suppressing KRAS-mutant PDAC and LUAD growth.

### Glutamine Antagonist DRP-104 Enhances RAS inhibitor Anti-tumor Activity

It has been shown that GLSi exhibited limited efficacy in vivo, which may result from cancer cells compensating for glutamate depletion following prolonged GLSi treatment (84). Thus, we hypothesized that an alternative approach targeting global glutamine metabolism may be necessary to achieve a pronounced anti-tumor effect in vivo. Recently, DRP-104, a prodrug of a potent glutamine antagonist 6-diazo-5-oxo-l-norleucine (DON), was developed to globally target glutamine-utilizing enzyme and found selective in malignant tumors (85). We and others have shown that DRP-104 can effectively and sustainably suppress *KEAP1* mutant LUAD and PDAC tumor growth with tolerable toxicity (86,87). Moreover, the drug is currently under phase 2 clinical evaluation in *KEAP1* or *NFE2L2* altered NSCLC patients (NCT07249372) and a phase 1/2 clinical trial assessing the combination with the anti-PD-L1 inhibitor durvalumab in fibrolamellar hepatocellular carcinoma (NCT06027086). Therefore, we hypothesized that DRP-104 treatment could sensitize tumors to KRAS inhibition.

We first tested the combination of RASi and DRP-104 in two WT and four *KRAS*-mutant PDAC patient-derived organoids (Supplementary Fig. S6A). Among these organoid cultures, PANFR0185 harbors WT *NRF2* amplification (copy number >5) and PANFR0165 contains the *NRF2*^D29H^ mutation which impairs interaction with KEAP1(88). RASi demonstrated potent inhibition in organoid growth in all KRAS mutant but not WT organoids (Fig. 6A and 6B; Supplementary Fig. S6B). All six organoids were sensitive to DRP-104, with PANFR0185 showing especially high sensitivity to DRP-104 (Supplementary Fig. S6C). Treatment with both drugs enhanced the growth suppression of these organoids compared with monotherapy. These observations support the therapeutic potential of co-targeting KRAS and glutamine metabolism in the setting of NRF2 activation.

**Figure 6.**
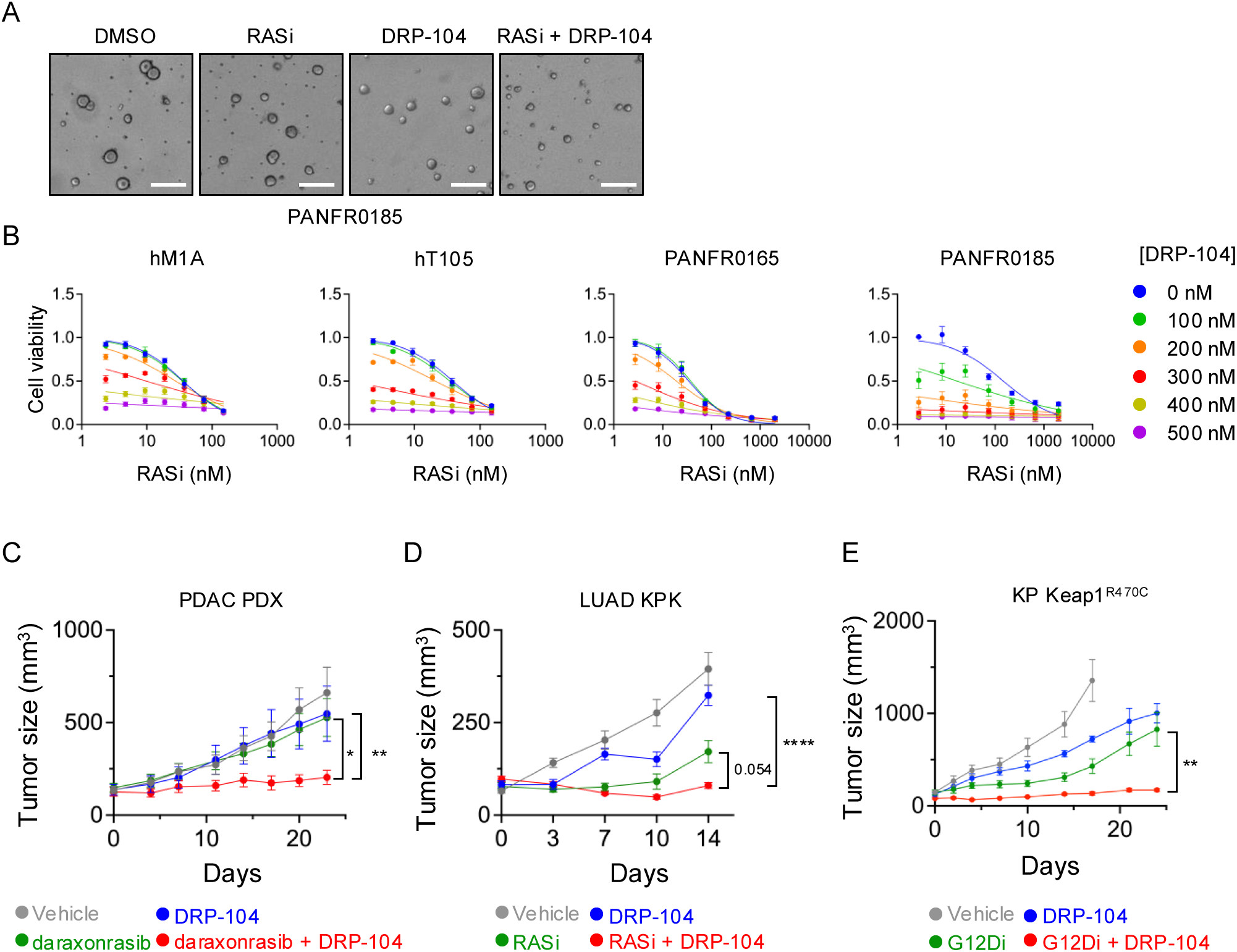
The combination of glutamine antagonist DRP-104 with RAS inhibitors effectively suppresses KRAS-mutant PDAC and LUAD tumor growth. **A,** Representative images of PANFR0185 organoids cultured in medium containing DMSO, RASi (24.7 nM), DRP-104 (400 nM), and the combination of both drugs (RASi + DRP-104). The scale bars denote 50 μm. **B,** PDAC patient derived organoids were treated with indicated concentrations of RASi and DRP-104 for 5 days, and cell viability was determined by 3D CellTiter Glo. Normalized cell viability compared to the control conditions, which were normalized to 1.0, were plotted, and error bars denoted standard error of at least three independent biological replicates. **C,** PANFR0185 organoids were subcutaneously injected into mice to induce PDAC patient-derived xenografts (PDX). Mice were randomly assigned to groups treated with vehicle, 3 mg/kg daraxonrasib, 3 mg/kg DRP-104, or the combination of both drugs, and tumor sizes were monitored for 23 days. **D,** KPK lung cancer cells were subcutaneously injected into mice to induce allografts. Mice were randomly assigned to groups treated with vehicle, 10 mg/kg RASi, 3 mg/kg DRP-104, or the combination of both drugs, and tumor sizes were monitored for 14 days. **E,** KP lung cancer cells expressing Keap1^R470C^ were subcutaneously implanted in mice to induce xenograft. Mice were treated with vehicle, 10 mg/kg G12Di, 3 mg/kg DRP-104, and the combination of the two drugs, and the tumor sizes were monitored for 24 days or until tumors reached the sacrificing endpoint.

Finally, we extended the analyses of this combination *in vivo* in KEAP1 or NRF2 altered models. First, we utilized the PANFR0185 (*KRAS*^G12R^/*NRF2*^Amp^) organoid culture to induce xenograft tumors in mice, and we evaluated the anti-tumor effect of concurrent treatment with G12Di, clinical candidate analog of RMC-7977 (RASi), the RAS(ON) multi-inhibitor daraxonrasib (RMC-6236) and DRP-104. The dosages and schedules were selected based on previously established protocols demonstrating potent anti-tumor activity and high therapeutic selectivity (86,89). Monotherapy with either DRP-104 or daraxonrasib exhibited limited anti-tumor activity. In contrast, the combination of DRP-104 and daraxonrasib significantly suppressed the tumor growth (Fig. 6C). Consistent with these findings, combining RASi with DRP-104 exhibited potent anti-tumor effect in mice bearing lung KPK subcutaneous tumors (Fig. 6D). Importantly, both experiments demonstrated limited toxicity profiles for either monotherapy or the combination, as there was no significant weight loss observed across different conditions (Supplementary Fig. S6D and S6E).

We then examined the combination of G12Di and DRP-104 in lung tumors induced by subcutaneous implantation of the *Kras*^G12D^/*p53*^Δ/Δ^/*Keap1*^R470C^ mouse LUAD cells (44). Consistent with the RASi studies, the treatments were tolerable and did not cause significant weight loss in mice (Supplementary Fig. S6F). Although treating mice with G12Di or DRP-104 alone was able to reduce the rate of tumor growth, the combination caused near-complete suppression of tumor growth (Fig. 6E). Collectively, these results support the potential of combination therapy targeting KRAS and glutamine metabolism in inhibiting KRAS-mutant, KEAP1-mutant or NRF2 activated pancreatic and lung cancers.

## DISCUSSION

KEAP1 mutations have been associated with poor patient responses to KRAS^G12C^ inhibitors, the molecular mechanisms underlying this resistance remain unclear (16,17,23,39). NRF2 overexpression caused by KEAP1 loss-of-function has been linked to reduced sensitivity to KRAS^G12C^ inhibitors in NSCLC patients (39). While *KEAP1* mutation is not commonly found in PDAC, the increased expression of NRF2 at advanced stages of PDAC highlights the importance of potential NRF2-mediated resistance (29,31). Here, we elucidated the mechanism of resistance to KRAS inhibition mediated by the KEAP1-NRF2 pathway to identify pharmacologic approaches to enhance the efficacy of KRAS-targeted therapy.

We determined that *KEAP1* depletion conferred NRF2-dependent resistance to KRAS inhibitors and established a KEAP1-dependent transcriptional program in PDAC. Our *KEAP1*^KO^ UP/DN signature differed significantly from established KEAP1/NRF2 gene sets, in part because of genetic (KRAS mutation) and cell type differences. This heterogeneity between the signatures is likely due to the diverse approaches to activate NRF2 (small molecule inducers, KEAP1 silencing, NRF2 overexpression), the methodologies (RNA-seq, ChIP-seq, or literature curation), different cancer or cell line models (osteosarcoma, lymphoblastoid, lung cancer patients, HEK293T, and PDAC), and distinct gene-selection criteria in generating each signature. There is a core of nine NRF2-regulated genes (*GCLC, GCLM, ME1, PIR, GSR, SLC7A11, NQO1, TXNRD1,* and *SRXN1*) that are shared across nearly all of these gene sets. Overall, our *KEAP1*^KO^ UP/DN signature, along with others, reflects the transcriptomic alterations under different contexts, and the convergence on core NRF2-regulated targets among them implicates that fundamental transcriptional program of NRF2 was generally captured.

The resistance to KRAS inhibitors has been linked to activation of KRAS-MYC and YAP/TAZ-TEAD signaling pathways (12,17,20,22). Here, we compared the upregulated transcriptome associated with KEAP1-deficient, KRAS-, MYC-, and TEAD-dependent programs (45,62,63). The KRAS-, MYC-, and TEAD-dependent, upregulated transcriptome are interconnected and regulate many essential genes promoting tumor progression, based on DepMap gene essentiality scores. However, the KEAP1-deficient transcriptome was found distinct from the other three and contains primarily non-essential genes. These findings are consistent with the nature of KEAP1-NRF2 pathway, which activates cytoprotective genes primarily under stress conditions (24). The dysregulation of KRAS, MYC, and TEAD has been established as drivers to cancer progression, agreeing with these highly essential genes detected under these axes (3,90,91). Notably, the unique profile of *KEAP1*^KO^ transcriptome highlights the heterogeneity of resistance mechanisms to KRAS inhibition, implicating the importance of tailored therapeutics to address patients with tumor relapsed after the KRAS inhibitor treatment under different genetic contexts.

Moreover, we determined a 200-gene PDAC KEAP1-deficient (PKD) signature comprising commonly and significantly upregulated genes upon KEAP1 loss. By cross referencing the ChIP-seq results from the ENCODE database as well as HOMER motif analysis, genes that are directly regulated by NRF2 consist of 50% of the signature. Collectively, these findings further verify the hypothesis that NRF2 is the major driver of the resistance to KRAS inhibition promoted by KEAP1 knockout. Importantly, the PKD signature stands out to be highly correlated to the poor drug response to KRAS inhibitors in both KRAS-mutant PDAC and lung cancers. Future integration of more extensive clinical sequencing datasets will be critical to validate our findings and to establish the PKD signature as a predictive biomarker in the responses of KRAS inhibitors in patients.

Our RNA-seq analysis also revealed an increased dependence of KEAP1-deficient cells on glutamine metabolism, driven in part by NRF2-induced upregulation of SLC7A11. We exploited the anti-tumor effects of combining the clinical candidate, glutamine antagonist DRP-104 with inhibitors targeting RAS, and we demonstrated that this combination strongly suppressed proliferation in PDAC patient-derived organoids, especially those with NRF2 gain-of-function alterations. In addition, this combination also potently inhibited KEAP1/NRF2-altered PDAC or LUAD tumor growth *in vivo* with no overt toxicity. These results support the potential of future clinical evaluation of combining DRP-104 with a RAS inhibitor in targeting KEAP1-mutant lung cancer or NRF2-overexpressing PDAC.

In conclusion, our study comprehensively characterized the KRAS inhibitor-resistant transcriptome induced by KEAP1 depletion, and we derived a PDAC KEAP1-deficient (PKD) gene signature from the RNA-seq transcriptomic analysis. The transcriptome is not only highly driven by NRF2 activation but also strongly associated with resistance to KRAS inhibitors. This mode of resistance is largely distinct from established mechanisms driven by KRAS, MYC, or TEAD. Building upon the transcriptomic profiling, we identified a drug combination co-targeting KRAS and glutamine metabolism and showed that this treatment gave rise to potent anti-tumor activities. These findings highlight mechanistic insight into KEAP1-NRF2-mediated resistance and provide a rationale for future clinical evaluation of combinations involving KRAS inhibitors and the glutamine antagonist in PDAC and NSCLC.

## METHODS

### Cell Line, Spheroid, and Organoid Cultures

RASless MEFs stably expressing exogenous KRAS or BRAF were provided by the National Cancer Institute (NCI) RAS Initiative and established as described previously (36). Pa01C (RRID: CVCL_E303), Pa14C (RRID: CVCL_1638), and Pa16C (RRID: CVCL_1639) were provided by Dr. Anirban Maitra (MD Anderson Cancer Center). HEK293T (RRID: CVCL_0063), PANC-1 (RRID: CVCL_0480), SW1990 (RRID: CVCL_1723), AsPC-1 (RRID: CVCL_0152), HPAC (RRID: CVCL_3517), HPAF-II (RRID: CVCL_0313), SU.86.86 (RRID: CVCL_3881), NCI-H441 (RRID: CVCL_1561), A549 (CVCL_0023), NCI-H358 (RRID: CVCL_1559), and NCI-H2030 (RRID: CVCL_1517) were acquired from American Type Culture Collection (ATCC). KP LUAD cell lines were obtained from Dr. Tyler Jacks (Massachusetts Institute of Technology). Further generation of control KP cells, Keap1-KO (KPK), Keap1^WT^-, and Keap1^R470C^-expressing KPK cell lines were described previously (44,86). Cells were cultured in either DMEM (Thermo Fisher, #11995065) or RPMI-1640 (Thermo Fisher, #11875093) supplemented with 10% fetal bovine serum (FBS, Sigma #2442) and PennStrep (Thermo Fisher, #15140163). All cells were cultured in a humidified incubator under 37°C and 5% CO_2_.

To establish RAS inhibitor-resistant cells, PDAC cell lines were initially cultured in appropriate medium containing either RASi or G12Di at their GI_50_ concentrations. Once the cultures reached 90% confluency, they were then passaged at 2.0 × 10^6^ cells per dish and maintained in growth medium with inhibitor concentrations increased 50% from the previous passage. Cells would be continuously propagated until their sensitivity of the inhibitors reach 100 X of their original GI_50_ concentrations. Cells were then cultured in the medium with the final drug dosages for future experiments.

Spheroids were generated by seeding 3,000 – 10,000 cells in a well of a round-bottom, low adherent plate in appropriate medium. The plates were then subjected to 300 × g centrifugation for 3 minutes. The plates were incubated in a humidified incubator under 37°C and 5% CO_2_ for two days to allow spheroid formation. The spheroids were then subjected to future experiments.

Patient-derived organoids were obtained from Dr. David Tuevson (Cold Spring Harbor Laboratory, hM1A, hT105) and the Dana-Farber Cancer Institute (PANFR0165, PANFR0185, PANFR0088, PANFR0344). The organoid culture protocol was described previously (92). In brief, the organoids were maintained in a humidified incubator under 37°C and 5% CO_2_. The organoids were cultured in reduced growth factor Matrigel domes with compete human feeding medium (Advanced DMEM/F12 (Thermo Fisher, #12634028)–based WRN conditioned medium (L-WRN)(ATCC, CRL-3276), 100 ng/mL hFGF10 (Thermo Fisher # 100-26), 10 mM nicotinamide (Sigma #72340), 0.01 μM hGastrin I (TOCRIS #3006), 50 ng/mL hEGF (Thermo Fisher # AF-100-15), 500 nM A83-01 (TOCRIS #2939), 1× B27 supplement (Thermo Fisher, #A1486701), 10 mM HEPES (Thermo Fisher #15630130), and 0.01 μM GlutaMAX (all Thermo Fisher, #35050079). All the patient-derived materials were obtained from patients with written consents. All the patient studies were conducted in accordance with the Declaration of Helsinki and approved by the Institutional Review Board.

### CRISPR-Cas9 Knockout

sgRNA (KEAP1-1, CTTGTGGGCCATGAACTGGG; KEAP1-2, TGTGTCCTCCACGTCATGAA; eGFP, GGGCGAGGAGCTGTTCACCG) (30) targeting *KEAP1* and *eGFP* were reported previously and cloned into LentiCRISPRv2 (Addgene #52961, RRID:Addgene_52961) or TLCV2 (Addgene #87360, RRID:Addgene_87360) (37). sgRNA (Keap1,CACCGCCGAAGCCCGTTGGTGAACA, AAACTGTTCACCAACGGGCTTCGGC; Neomycin resistant gene, CCTCGGAACAAGATGGATTGCACGC) for targeting *Keap1* or Neomycin resistant gene were cloned into lentiCRISPRv2 puro (Addgene #98290, RRID: Addgene_98280). pLenti-DECKO-Cas9-Blast construct cloned with sgRNA targeting NRF2 (GGAGTAGTTGGCAGATCCAC) were provided from Dr. M. Benjamin Major (Washington University at St. Louis). sgGFP-Cas9-Blast (PXPR007 sgGFP-1, Addgene #74960, RRID: Addgene_74960) construct was acquired from Addgene. *Keap1*^R470C^ or *Keap1*^WT^ was cloned into a Gibson compatible lentiviral construct as described previously (44).

Lentivirus was used to transduce CRISPR-Cas9 system into cells and was generated by transfecting HEK-293T cells with the cloned construct, psPAX2 (Addgene #12260, RRID: Addgene_12260), and pMD2.G (Addgene #12259, RRID: Addgene_12259) via FuGene 6 (Promega #E2691). HEK293T cells were incubated with the transfecting agents for 24 hours, and the medium was replaced with fresh medium supplemented with 10% FBS. After 48 hours, the conditioned medium was collected, filtered using a 0.45 μm filter, and stored at a -80°C freezer for future use.

For virus transduction, cells were cultured in the appropriate medium with the addition of the conditioned medium containing desired lentivirus and 8.0 μg/mL polybrene. The medium was replaced with fresh medium after 24 hours incubation. After an additional day, cells were then selected using 2.0 – 3.0 μg/mL puromycin (Thermo Fisher # A1113803) for 2 days, or 7.5 mg/mL blasticidin (Corning # 30-100-RB) for 7 days. For inducible CRISPR knockdown using TLCV2 construct, knockout was induced by culturing cells with 2.0 – 4.0 μg/mL doxycycline (Sigma #D5207) for at least 5 days. Cells were then subjected to further western blot, proliferation, apoptosis, spheroid, metabolomic, and RNA sequencing analysis.

### CRISPR-Cas9 Screen

A single-guide RNA (sgRNA) library targeting 2,240 genes was generated and provided by Dr. Kris Wood (Duke University) (37). PANC-1 cells were transduced with this library at multiplicity of infection (MOI) of 0.2 via lentivirus, followed by puromycin selection (3.0 μg/mL) for 2 days. Cells were harvested immediately after selection (day 0) and after 7 additional days in culture (day 7). Subsequently, cells were split into two groups: one treated with 10 nM G12Di and the other with DMSO. Both groups were continuously cultured under the indicated conditions for 4 weeks before harvested (day 35). Genomic DNA from samples collected at days 0, 7, and 35 was extracted using DNeasy Blood & Tissue Kit (Qiagen # 69504) and subjected to next generation sequencing (NGS). MAGeCK algorithm was used to determine the enrichment of sgRNA for each gene within the library(93).

### In Vitro Proliferation Assays

Five hundred – 3,000 cells were seeded in a well of a 96-well plate. Cells were treated with MRTX1133 (Selleckchem #E1051, or Wuxi Network Lab), RMC-7977 (MedChemExpress # HY-156498), AI-1 (Sigma # 492041), CDDO-Me (Sigma # SMB00376), CB-839/GLSi (Selleckchem #S7655), and DRP-104 (Selleckchem #E1203, or Wuxi Network Lab), indicated conditions and incubated for 5 days. Cell viability was measured using Calcein-AM, Hoechst or Cell TiterGlo cell viability assay. For Calcein-AM or Hoechst assay, cell culture medium was aspirated followed by the addition of 50 μL, 0.50 ng/μL Calcein-AM (Thermo Fisher #C34852) or 2 mM Hoechst (Thermo Fisher # H1398) dissolved in phosphate-buffered saline (PBS) to each well of the 96-well plates and incubated in 37°C for 15 minutes.

After incubation, the fluorescence images were taken using a plate reader (SpectraMax i3X MiniMax 300 Spectrometer, Molecular Devices; or Cytation 1 Cell Imaging Multi-Mode Reader, BioTek) at 465 nm excitation and 541 nm emission for Calcein AM fluorescence, or at 360 nm excitation and 450 nm emission for Hoechst. The individual cell numbers were counted by fluorescence images.

For CellTiter cell viability assay, CellTiter Luminescent cell viability assay (Promega #G7570) was used for 2D cell culture, and CellTiter-Glo 3D cell viability Assay (Promega #G9683) was performed for 3D spheroid or organoid cultures per manufacturer’s protocol. In brief, a volume of CellTiter reagents equivalent to the existing volume of culture medium in the well was added to the well containing cells or organoids and was mixed by vigorous pipetting for at least 20 times. The plate was then incubated at room temperature for 30 minutes to allow the luminescent signal developed. After the incubation, the solution was transferred to opaque, white plates and luminescent signal was quantified by a plate reader.

### NMR Metabolomic Analysis

One x 10^5^ cells were seeded on a 12-well plate and settled overnight. To start the experiment, the medium was aspirated, washed by PBS, and replaced by 750 μL of fresh low-glucose DMEM (Thermo Fisher #11885-084) or RPMI supplemented with 10% FBS for 24 hours. The conditioned medium was collected and filtered using a 10 KDa molecular weight cut off Amicon Ultra centrifugal unit for 30 minutes. Four hundred μL flow-through conditioned medium were then mixed with 200 μL of standard solution containing 6 mM trimethylsilylpropanoic acid (TSP) and 3 mM imidazole prepared in 99% D2O. The NMR spectrums were collected using a Bruker Advance III 700 mHz Spectrometer. The metabolites were determined and quantified using Chenomx.

### Western Blot Analyses

Cells were lysed by lysis buffer (20 mM pH 7.4 Tris-HCl, 150 mM NaCl, 2 mM EDTA, and 1% Triton X-100), and the total protein concentrations of lysates were quantified by Bradford Assay. Equal amounts of cell lysates (based on total protein mass) were loaded and resolved on SDS-PAGE gels, transferred to PVDF membranes, and blocked by 5.0% bovine serum albumin (BSA) in TBST buffer (20 mM Tris, 2.7 mM KCl, 140 mM NaCl, and 0.05% Tween-20). The membrane was incubated with the primary antibody with 1000X dilution in TBST in 4°C overnight. Subsequently, the membrane was incubated with the HRP-conjugated secondary antibody with 5000X dilution in TBST in room temperature for 1 hour. The membrane was then washed, incubated with ECL substrate, and imaged using a Bio-Rad ChemiDoc MP imager. The vendors, catalog numbers, and RRIDs of antibodies used in this study are documented in the Supplementary Table 4.

### siRNA Knockdown

One x 10^5^ cells were first seeded in a well of six-well plates and incubated in the incubator overnight. siRNAs targeting SLC7A11 (Horizon Discovery, siSLC7A11-1, UUGUAGAGCUUGCGUAGAA, # J-007612-09; siSLC7A11-2, CGGCAAACUUAUUGGGUCU, # J-007612-10) or siRNA with non-specific sequence (Thermo Fisher #4390844) were mixed with OptiMEM (Thermo Fisher #31985088) and lipofectamine RNAiMAX (Thermo Fisher #13778150) and incubated in room temperature for 15 minutes. The siRNA mixture was then introduced to the cells and incubated for 24 hours. Then, medium containing siRNA was removed, and cells were subjected to western blot or proliferation assay for further analysis.

### Cell Death Assay

Seven thousand - 10,000 PDAC cells were seeded in a 96-well plate with appropriate medium supplemented with CellTox Green Express (Promega #G8731). Cells were incubated overnight, and the baseline fluorescence signals were read using a plate reader at 485 nm excitation and 535 nm emission. Cells were then treated with indicated concentrations of G12Di. After 3 days, the final fluorescence signals were read. The normalized apoptosis flux is defined as:

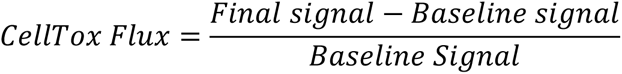

### Oxidative Stress Assay

The oxidative stress assay was determined using CellROX™ Green (Thermo Fisher # C10444) per manufacturer’s protocol. In brief, 50,000 to 100,000 cells were seeded in petri dishes or six-well plates and allowed to adhere overnight. Cells were then treated with the indicated concentrations of drugs for 72 hours. Following the treatment, cells were stained by 2.0 µM CellROX Green reagent or DMSO for 1 hour. Cells were then washed twice with PBS, harvested, and pelleted by centrifugation at 300 × g for 5 minutes. Cell pellets were resuspended in DMEM and analyzed by flow cytometry using a CytoFLEX S flow cytometer (Beckman Coulter, USA) with 488 nm excitation and 525 nm emission. A minimum of 20,000 cells were recorded to quantify ROS levels.

### Cell Cycle Assay

Cells were seeded in petri dishes and allowed to adhere overnight. Cells were then treated with the indicated conditions of drugs for 72 hours. After the treatment, culture media and cells were collected, washed by PBS, and pelleted under 300 x g centrifugation for 5 minutes. Cell pellets were resuspended in 400 µL of ice-cold PBS. To fix the cells, 800 µL of ice-cold 100% ethanol was added and mixed gently, and the samples were incubated at 4°C for two hours to ensure complete fixation. Fixed cells were centrifuged to remove ethanol and incubated with FxCycle™ PI/RNase Staining Solution (Thermo Fisher # F10797) for 15-30 minutes at room temperature, protected from light. The stained cells were analyzed by a CytoFLEX S flow cytometry with 561 nm excitation and 610 nm emission. A minimum of 20,000 cells were recorded. Cell debris was excluded by SSC-A versus FSC-A gating, and single cells population were identified by FSC-H versus FSC-A gate. A histogram of propidium iodide signal was generated from the singlet cell population, and the G1, S, and G2/M phases of the cell cycle were determined using FCS Express 7 Research (De Novo Software).

### Glutaminase Activity Assay

Glutaminase activity was determined by a two-step reaction coupling NAD^+^ reduction to glutamine hydrolysis using the mitochondria fraction of cells, as described previously (94,95). For mitochondrial isolation, cells were cultured in petri dish plates until they reached 90-100% confluency. The cells were washed with PBS, harvested, pelleted by centrifugation (600 × g) for 5 minutes, and resuspended in 1000 μL ice-cold isolation buffer (10 mM pH 7.4 Tris-MOPS, 1 mM EGTA, 10 mM sucrose). The cells were homogenized using a Dounce homogenizer in ice and centrifuged at 600 × g for 10 minutes at 4°C. The supernatant was collected and centrifuged at 7,000 × g for 10 minutes at 4°C, and the resulting pellet containing mitochondria was washed with isolation buffer once and resuspended in isolation buffer.

To determine the glutaminase activity, 10 μg mitochondrial fractions were brought to a final volume of 10 μL by the isolation buffer containing GLSi at the indicated concentrations or DMSO and incubated at 37 °C for 10 minutes. Then, 105 μL of the first reaction mixture (65 mM pH 8.6 Tris-acetate, 0.25 mM EDTA, 75 mM NaCl, 17.4 mM L-glutamine, 130 mM K_2_HPO_4_) were added to initiate the reaction of glutamine to glutamate at 37°C for 30. The first reaction was quenched by the addition of 10 uL of 3.00 M HCl on ice. Subsequently, 20 μL of the first reaction mixture was then mixed with 200 μL of the second reaction mixture (130 mM pH 9.4 tris-HCl, 0.36 mM ADP, 1.80 mM NAD^+^, 1 unit of glutamate dehydrogenase (Sigma #10197734001)) and incubated at room temperature for 45 minutes to convert glutamate to 2 α-ketoglutarate, coupled with NAD+ to NADH conversion. The absorbance of NADH for the final reaction mixture was measured at 340 nm to determine the glutaminase activity

### Clonogenic Growth Assay

Five thousand – 10,000 cells were seeded in a well of a six-well plate and settled overnight. Cells were treated with indicated concentrations of G12Di, and fresh medium with G12Di were replenished every 4-5 days. After 14 days, cells were washed by PBS twice and fixed by 3.7% formaldehyde. The fixed cells were then stained by 0.5% crystal violet for 10 minutes, washed by PBS three times, and imaged using a Typhoon FLA9500 biomolecular imager.

### RNA Sequencing

For PDAC studies, 10^5^ cells were seeded in 6-cm petri dishes. The cells were treated with GI_50_ concentrations of G12Di or DMSO for 3 days. The cells were snap-frozen by liquid nitrogen, and the RNA was extracted using Qiagen RNeasy plus kit (Qiagen # 74134). The samples were frozen in -80°C and submitted to Novogene (USA) for poly A enrichment, directional mRNA library preparation using NEBNext^®^ Ultra™ II Directional RNA Library Prep Kit (New England Biolabs #E7760), and next generation sequencing using a Novaseq 6000 sequencer.

For KP lung cancer models, 10^3^ cells were seeded as spheroids in media containing KI696 and treated with G12Di for 4 days. The spheroids were harvested and snap-frozen, and RNA was extracted using a Purelink RNA mini kit (Invitrogen). mRNA was converted to cDNA by using a SMARTer PCR cDNA synthesis kit (Clonetech), followed by sequencing library preparation via Nextera XT DNA library preparation kit (Illumina). Samples were pooled at equimolar ratios, loaded on SP11 cycle flow cells, and sequenced on Illumina NovaSeq600.

The quality of the NGS was verified by FastQC (v0.12.1). The fastq files were aligned to human reference genome (GRCh38.p14 or GRCm38) by STAR (v2.7.11b). The raw counts of mRNA transcripts were quantified using featureCounts within Samtools (v1.22) package or RSEM (v1.3.3). The following analyses and packages were performed in R environment: differential expression analyses for each paired condition were done using DESeq2 (v1.42.1) or limma (v3.58.1); and the pathway enrichment analyses were conducted by fgsea (v1.28.0), clusterProfiler (v4.10.1), and Singscore (v1.22.0) (96). Motif analyses were defined from -1000 to +1000 nucleotides from the transcription start site (TSS) by HOMER (v5.1) (48).

### ChIP-seq Analysis

Optimal IDR thresholded peak files from NRF2/NFE2L2 chip-sequencing experiments were downloaded from ENCODE for HeLa, A549, IMR-90, and Hep G2 cell lines (Acc: ENCFF305KIK, ENCFF418TUX, ENCFF474PPT, ENCFF882YL0). Promoters were defined based on Hg38 Gencode (v30) (97) annotations and summarized to genes using the R package TxDB.Hsapiens.UCSC.hg38.knownGene (v3.21.0). Overlaps between gene promoters and peak files were conducted with the IRanges (v2.38.1) and GenomicRanges (v1.56.2) R packages (98). Gene promoters were prioritized based on frequency of peak occurrence across the four cell line experiments and annotated using BiomaRt (v2.50.3) (99).

### Network Analysis

Differential analysis were carried out by limma on RNA-seq data in *KEAP1*^KO^ PDAC cells, PDAC cells treated with siRNA targeting KRAS (KRAS UP) (45) or MYC (MYC UP) (62), or cells treated with TEAD inhibitors (GNE-7883, TEAD UP; GSE273175) (63). The log_2_FC is normalized by scaled adjusted P values using the following equation to generate weighted log_2_FC. Genes with expressions significantly upregulated (adj. *p* < 0.05, log_2_FC > 0) were filtered and subjected to network analysis. The network analysis was conducted using R packages tidygraph (v1.3.1) and igraph (v2.1.4), plotted by ggraph (v2.2.1).

### Cancer Gene dependency scores, Mutational Data, and RNA expression

The gene dependency scores, mutational data of cancer cell lines for PDAC and LUAD were acquired from Cancer Dependency Map (DepMap). The data were accessed via R package depmap (v1.16.0). The RNA expressions for TCGA-LUAD cohort were acquired from TCGA database.

### Mouse Study

All mouse experiments described in this study were approved by the Institutional Animal Care and Use Committee (IACUC) at New York University (NYU). Mice were housed according to IACUC guidelines in ventilated cages in a pathogen-free animal facility. For all mouse studies, ≥ 4 mice were used for each condition. All transplantation experiments were performed using C57BL/6 J WT (JAX strain 000664) or NOD SCID Gamma (NSG; JAX strain 005557 F) mice. For cell line-derived xenograft experiments, 500,000 cells suspended in 100 μl PBS were injected subcutaneously into the flanks of the experimental mice. For the PDAC patient-derived xenograft experiment, PDAC organoids cultures were homogenized and injected subcutaneously into NSG mice, and after tumor outgrowth, the mice were euthanized, the tumors excised, cut to 25 mm^3^ pieces and surgically implanted into naive NSG hosts. Tumors were measured with calipers, and volume was calculated based on 0.5 (length × width^2^). Mice were euthanized when tumors reached the maximum volume of 2,000 mm^3^ permitted by IACUC at NYU, or sooner if the tumors reached pre-determined endpoints.

For each experiment, once the tumors of a mouse treatment cohort reached 100 – 250 mm^3^, mice were randomized into respective treatment groups and dosed at the indicated conditions until experimental endpoint. G12Di was dosed at 10 mg/kg or vehicle (50 mM pH 5.0 sodium citrate + 10% captisol) by intraperitoneal injection (daily). Daraxonrasib (RMC-6236) (MedChemExpress # HY-148439) was dosed at 3 mg/kg or vehicle (10% DMSO, 20% PEG400, 10% Solutol HS 15, in water) by oral gavage (daily). RMC-7977/RASi was dosed at 10 mg/kg (same vehicle as daraxonrasib) by oral gavage (daily). DRP-104 was dosed at 3 mg/kg or vehicle (10% Tween 80, 10% ethanol, in 0.9% saline) by subcutaneous injection (five days on, two days off schedule).

### Statistical Analyses

Computational analyses were conducted using R (v4.3.2) and RStudio (v2024.9.0.375). All experiments are reported with at least three independent samples. Quantitative results are reported as mean ± standard error of the mean (SEM). Statistical analyses are performed using GraphPad Prism software, and significance is evaluated using Student’s t-tests, Wilcoxon test, or ANOVA. Significance thresholds are defined as follows: *p < 0.05, **p < 0.01, ***p < 0.001, and ****p < 0.0001.

## Data Availability

Additional data generated in this study are available from the corresponding author upon request.

## Authors’ Disclosures

C. A. Stalnecker has received consulting fees from Reactive Biosciences. K.L. Bryant has received research funding support from SpringWorks Therapeutics. A.D. Cox has consulted for Eli Lilly and Mirati Therapeutics, Inc., a Bristol Myers Squibb company. T. Papagiannakopoulos received funding from Pfizer Medical Education Group, Dracen Pharmaceuticals, Kymera Therapeutics, Bristol Myers Squibb, and Agios Pharmaceuticals not related to the submitted work. C.J. Der is a consultant/advisory board member for AskY Therapeutics, Cullgen, Deciphera Pharmaceuticals, Kestrel Therapeutics, Mirati Therapeutics, Reactive Biosciences, Revolution Medicines, and SHY Therapeutics. C.J. Der has received research funding support from Deciphera Pharmaceuticals, Mirati Therapeutics, Reactive Biosciences, Revolution Medicines, and SpringWorks Therapeutics. B.M. Wolpin has consulted for Agenus, BMS/Mirati, EcoR1 Capital, GRAIL, Harbinger Health, Ipsen, Lustgarten Foundation, Revolution Medicines, Tango Therapeutics, and Third Rock Ventures. B.M. Wolpin has received research funding from BMS/Celgene, BreakThrough Cancer, Eli Lilly, Harbinger Health, Lustgarten Foundation, NIH/NCI, Novartis, Pancreatic Cancer Action Network, Revolution Medicines, Servier/Agios, and Stand Up to Cancer. A.J. Aguirre. has consulted for Affini-T Therapeutics, AstraZeneca, Blueprint Medicines, Boehringer Ingelheim, Curie.Bio, Incyte, Kestrel Therapeutics, Merck & Co., Inc., Mirati Therapeutics Inc., Nimbus Therapeutics, Oncorus, Inc., Plexium, Quanta Therapeutics, Reactive Biosciences, Revolution Medicines, Riva Therapeutics, Servier Pharmaceuticals, Syros Pharmaceuticals, Taiho Pharmaceuticals, T-knife Therapeutics, Third Rock Ventures, and Ventus Therapeutics; holds equity in Riva Therapeutics and Kestrel Therapeutics. A.J. Aguirre. has research funding from Amgen, AstraZeneca, Boehringer Ingelheim, Bristol Myers Squibb, Deerfield, Inc., Eli Lilly, Mirati Therapeutics Inc., Novartis, Novo Ventures, Revolution Medicines. K.-K. Wong has consulted for and/or has received Grant/Research support from: Tango Therapeutics, Janssen Pharmaceuticals, Pfizer, Bristol Myers Squibb, Zentalis Pharmaceuticals, Blueprint Medicines, Takeda Pharmaceuticals, Mirati Therapeutics, Novartis, Genentech, Merus, Bridgebio Pharma, Xilio Therapeutics, Allerion Therapeutics, Boehringer Ingelheim, Cogent Therapeutics, Revolution Medicines and AstraZeneca. Other authors declare no competing interests.

## Authors’ Contributions

**W.-H. Chang:** Conceptualization, formal analysis, investigation, visualization, methodology, project administration, writing-original draft, and editing. **A. Vaughan:** Conceptualization, formal analysis, investigation, visualization, methodology, writing original draft, and editing. **A. G. Stamey:** Conceptualization, formal analysis, investigation, visualization, methodology, and writing-original draft. **M. Mancini:** Conceptualization, formal analysis, investigation, visualization, and methodology. **M. Hayashi**: Conceptualization, formal analysis, investigation, visualization, and methodology. **R. Yang:** Formal analysis and investigation. **R. Robb:** Formal analysis and investigation. **J.A. Klomp:** Formal analysis, investigation, and methodology. **A. M. Waters:** Conceptualization, formal analysis, investigation, and methodology. **A. Schaefer:** Formal analysis and investigation. **D. Andrussier:** Formal analysis and investigation., **K.L. Bryant:** Resources, supervision, and funding acquisition. **A.D. Cox:** Resources, supervision, editing, and funding acquisition. **F.M. Simabuco:** Resources, supervision, and funding acquisition. **K.-K. Wong:** Resources, supervision, and funding acquisition., **C.A. Stalnecker**: Formal analysis, investigation, methodology, writing-original draft, and editing. **T. Papagiannakopoulos:** Conceptualization, resources, supervision, funding acquisition, and editing. **C.J. Der:** Conceptualization, resources, supervision, funding acquisition, project administration, writing-original draft, and editing.

## Acknowledgments

W.-H. Chang was supported by a fellowship from the American Cancer Society (ACS) PF-23-1072348-01-CDP. R. Robb was supported by National Cancer Institute (NCI) T32CA071341 and F31CA284869. C.A. Stalnecker was supported by NCI T32CA009156 and F32CA232529. A.M. Waters was supported by ACS fellowship PF 18-061 and by NCI K22CA276632-01. K.L. Bryant was supported by Pancreatic Cancer Action Network (PanCAN)/AACR grant 15-70-25-BRYA, NCI P50CA196510 and R37CA251877, Department of Defense W81XWH2110693, and a Sky Foundation grant. Support was provided by grants to A.D. Cox and/or C.J. Der from the NCI; R01CA42978, P50CA196510, P50CA257911, U01CA199235, P01CA203657 and R35CA232113, to C.J. Der, from the Pancreatic Cancer Action Network/AACR (15-90-25-DER) and PanCAN 22-WG-DERB, and the Department of Defense (W81XWH2110692). T. Papagiannakopoulos was supported by NIH grants (R37CA222504, R01CA227649, R01CA283049, R01CA262562) and an American Cancer Society Research Scholar Grant (RSG-17-20001–TBE). A.J. Vaughan is supported by the NIH training grant (T32GM136542). K.-K. Wong was supported by a NIH grant (R01CA293718). A.J. Aguirre was supported by the Lustgarten Foundation, the Pancreatic Cancer Action Network, the Hale Center for Pancreatic Cancer Research, Break Through Cancer, NIH–NCI grants (P50CA127003, U01CA274276 and R01CA276268), and the Dana-Farber Cancer Institute Hale Center for Pancreatic Cancer Research. F.M. Simabuco was supported by the São Paulo Research Foundation (FAPESP (2018/14818-9) and a National Council for Scientific and Technological Development (CNPq) fellowship (303869/2021-6). The NMR studies was performed at The UNC Biomolecular NMR Laboratory, and it was supported by an NCI grant (P30CA016086). We thank Stuart Parnham for technical support on the NMR studies.

## Supplementary Figures

**Supplementary Figure S1.**
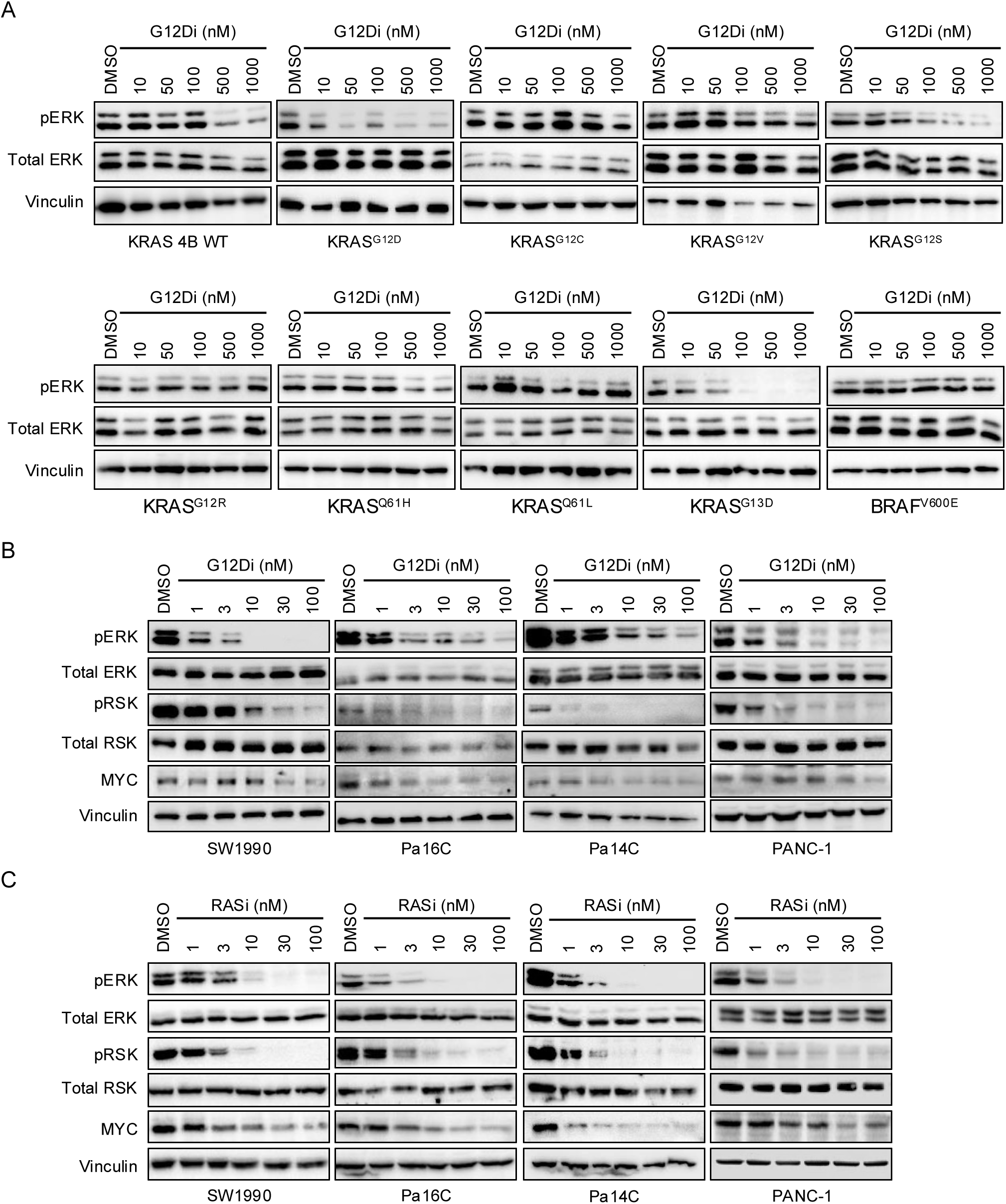
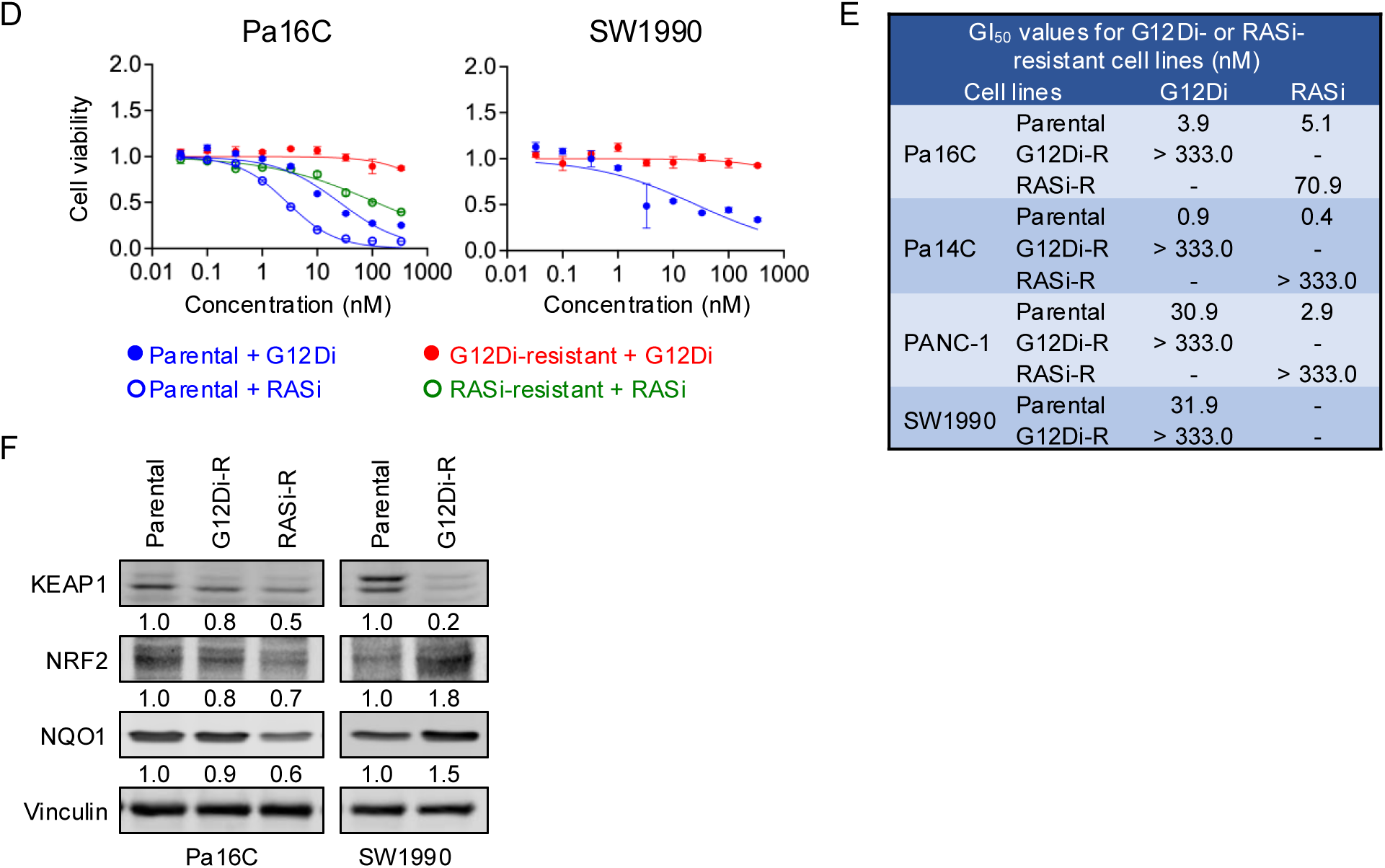
**A,** Representative western blots probing phospho-ERK 1/2 (pERK), total ERK 1/2 (ERK), and vinculin in RASless MEFs carrying different alleles of mutant KRAS or BRAF (indicated below the blots) and treated with G12Di (0 -1000 nM) for 4 hours. **B,** Western blot analyses in PDAC cell lines treated with G12Di for 4 hours. **C,** Representative western blots probing phospho-ERK1/2 (pERK), total ERK1/2 (ERK), phospho-RSK, total RSK, MYC, and vinculin in PDAC cell lines (SW1990, Pa16C, Pa14C, and PANC-1) treated with RASi (0 - 100 nM) for 4 hours. **D,** Resistant or parental PDAC cell lines were treated with G12Di (0 - 333 nM) or RASi (0 - 333 nM) for 5 days, and cell viability was determined by the addition of Hoechst. Normalized cell counts compared to the control conditions, which were normalized to 1.0, were plotted, and error bars denoted standard error of at least three independent biological replicates. **E,** Table of GI_50_ of G12Di and RASi on parental, G12Di-resistant (G12Di-R), or RASi-resistant (RASi-R) PDAC cells (SW1990, Pa16C, Pa14C, and PANC-1). **F,** Representative western blots probing KEAP1, NRF2, NQO1, and vinculin in Pa16C or SW1990 cells which are resistant to G12Di (G12Di-R), RASi (RASi-R), or their parental counterpart. The relative levels of KEAP1, NRF2, and NQO1 expressions are denoted underneath the designated blot images.

**Supplementary Figure S2.**
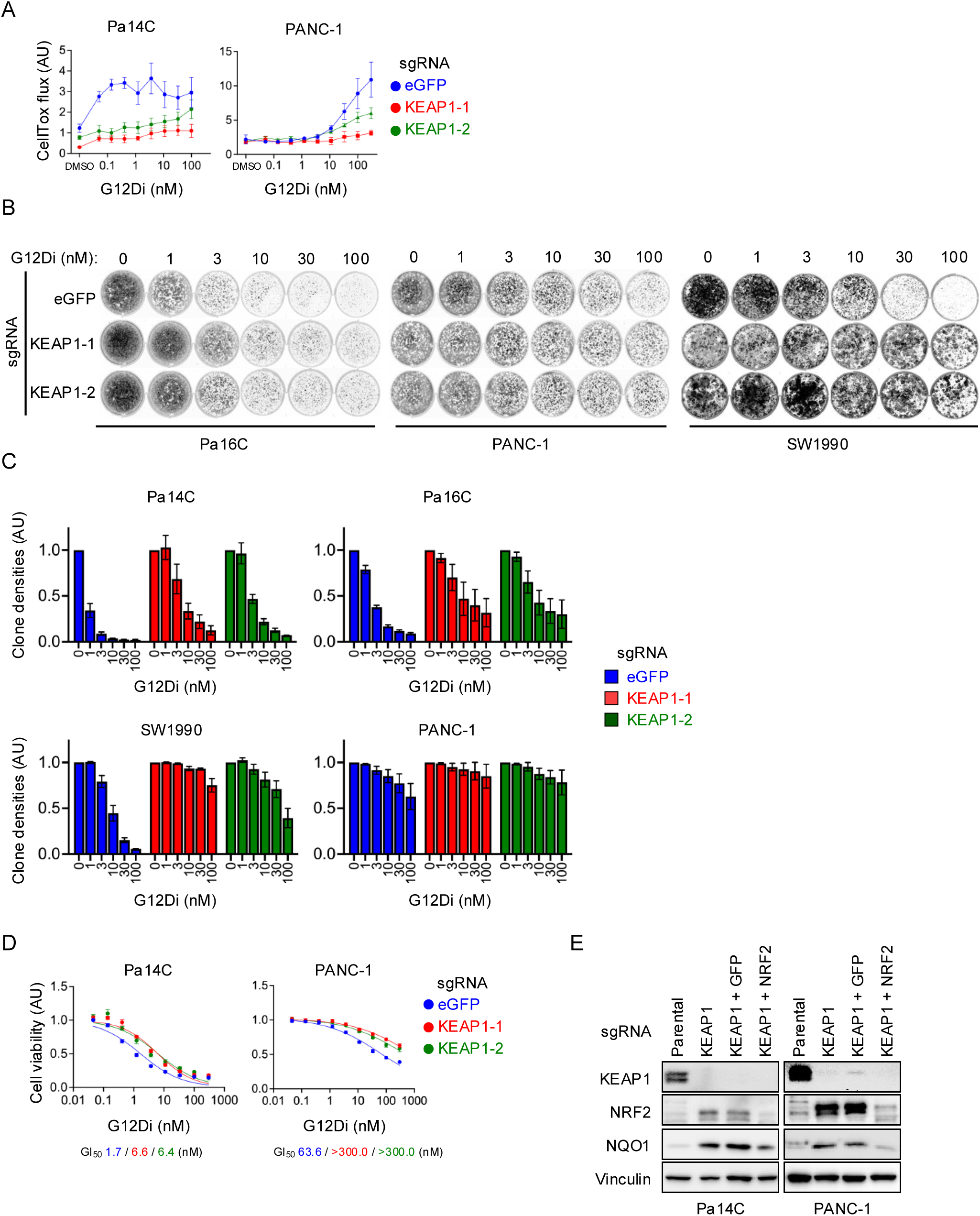
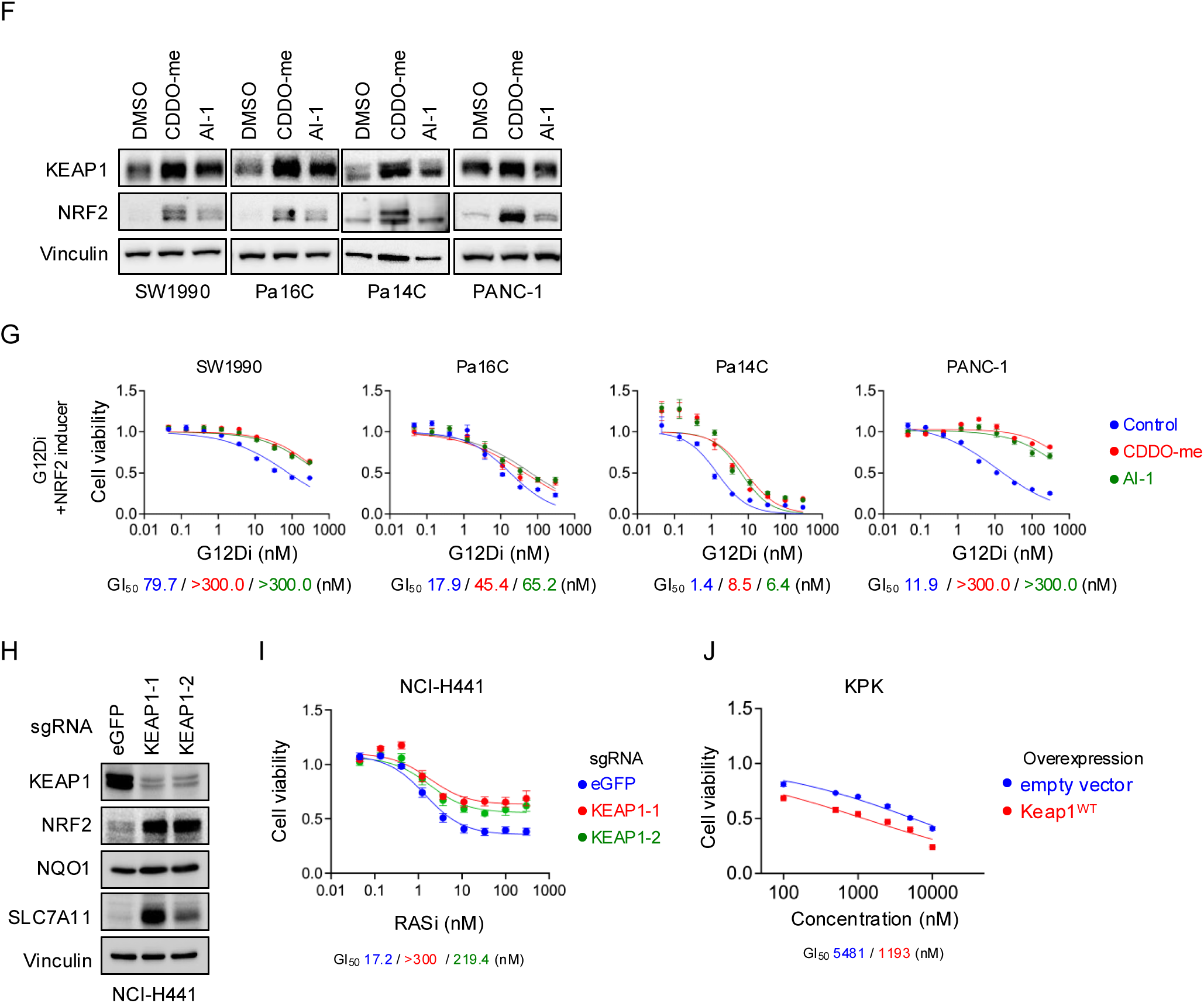
**A,** PDAC cell lines (PANC-1 and Pa14C) with *KEAP1* knockout (KEAP1-1, red; and KEAP1-2, green), or *eGFP* knockout controls (eGFP, blue), were treated with G12Di (0 – 300 nM) for 3 days, and the levels of cell death were assessed using CellTox^TM^ Green cytotoxicity assay (CellTox flux in arbitrary unit). **B,** Representative 2D clonogenic growth assays show long-term cell proliferation of SW1990, Pa16C, and PANC-1 cells upon knockout of *KEAP1* (KEAP1-1 and KEAP1-2) or eGFP. Cells were cultured with G12Di (0 – 100 nM) for 14 days, and they were stained at the end of the experiment with crystal violet and imaged using a Typhoon FLA9500 biomolecular imager. **C,** Quantification of cell densities of PDAC cells from the clonogenic growth assays in Figure 2D and Supplementary Figure S2B. Colony densities compared to the DMSO (0 nM G12Di) in each condition were normalized to 1.0, and error bars denoted standard error of at least three independent biological replicates. **D,** PDAC cell lines (PANC-1 and Pa14C) with *KEAP1* knockout (KEAP1-1, red; and KEAP1-2, green), or control cells (eGFP, blue), were grown using ultra-low attachment plates to form spheroid and treated with G12Di (0 – 300 nM) for 5 days, and cell viability was determined by 3D CellTiter-Glo at the end of experiment. Normalized cell counts compared to the control conditions, which were normalized to 1.0, are plotted, and error bars denoted standard error of at least three independent biological replicates. **E,** Representative western blots probing KEAP1, NRF2, NQO1, and vinculin in Pa14C and PANC-1 parental cells, cells with *KEAP1* knockout (KEAP1), *KEAP1* and *GFP* double knockout (KEAP1 + GFP), as well as *KEAP1* and *NRF2* double knockout (KEAP1 + NRF2). **F,** Representative western blots probing KEAP1, NRF2, and vinculin in PDAC cell lines treated with 50 nM CDDO-me, 5 µM AI-1, or DMSO as controls, for 24 hours. **G,** PDAC cell lines (SW1990, Pa16C, Pa14C, and PANC-1) were treated with G12Di (0.00 – 300 nM) and one of the NRF2 inducers, AI-1 (5.0 µM, red) or CDDO-me (50.0 nM, green), or DMSO (blue) as controls, for 5 days, and cell viability was determined by the addition of Calcein-AM assay. Normalized cell counts compared to the control conditions, which were normalized to 1.0, were plotted, and error bars denoted standard error of at least three independent biological replicates. Concentrations of G12Di that inhibit 50% viability were defined as GI_50_ and are shown in the corresponding colors underneath each plot. **H,** Representative western blots probing KEAP1, NRF2, NQO1, SLC7A11, and vinculin in H441 NSCLC cells with *KEAP1* knockout (KEAP1-1 and KEAP1-2) or control *eGFP* knockout (eGFP). **I,** H441 NSCLC cells upon *KEAP1* knockout (KEAP1-1, red; and KEAP1-2, green), or *eGFP* knockout controls (eGFP, blue), were treated with RASi (0.00 – 300 nM) for 5 days, and cell viability was determined by the addition of Calcein-AM assay. Normalized cell counts compared to the control conditions, which were normalized to 1.0, were plotted, and error bars denoted standard error of at least three independent biological replicates. Concentrations of RASi that inhibit 50% viability were defined as GI_50_ and are shown in the corresponding colors underneath each plot. **J,** KPK cells ectopically expressing Keap1^WT^ or an empty vector were treated with G12Di (0 – 10.0 μM) for 3 days, and cell viability was determined by 2D CellTiter Glo assay. Normalized cell counts compared to the control conditions, which were normalized to 1.0, were plotted, and error bars denoted standard error of at least three independent biological replicates. Concentrations of G12Di that inhibit 50% viability were defined as GI_50_ and are shown in the corresponding colors underneath each plot.

**Supplementary Figure S3.**
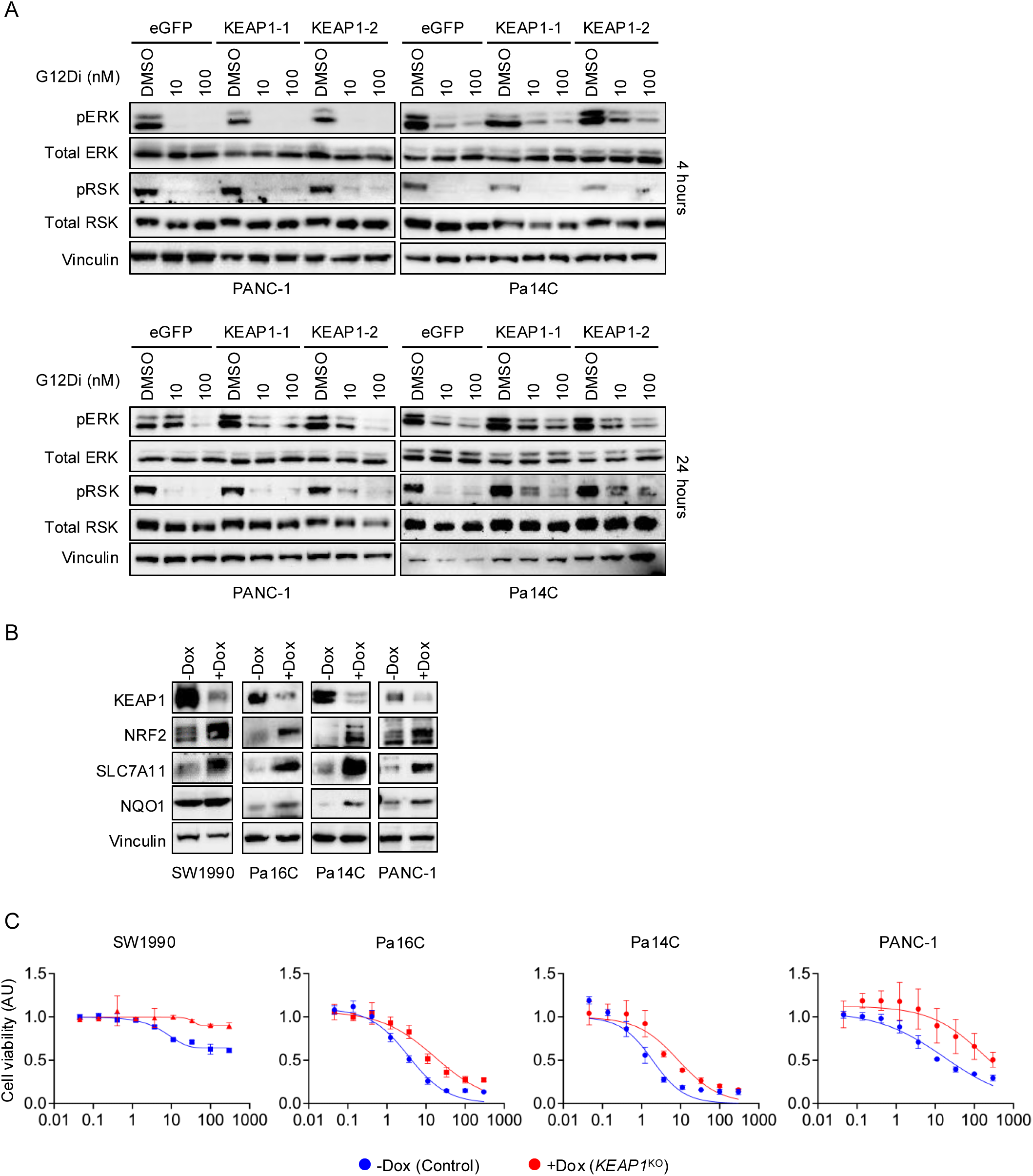
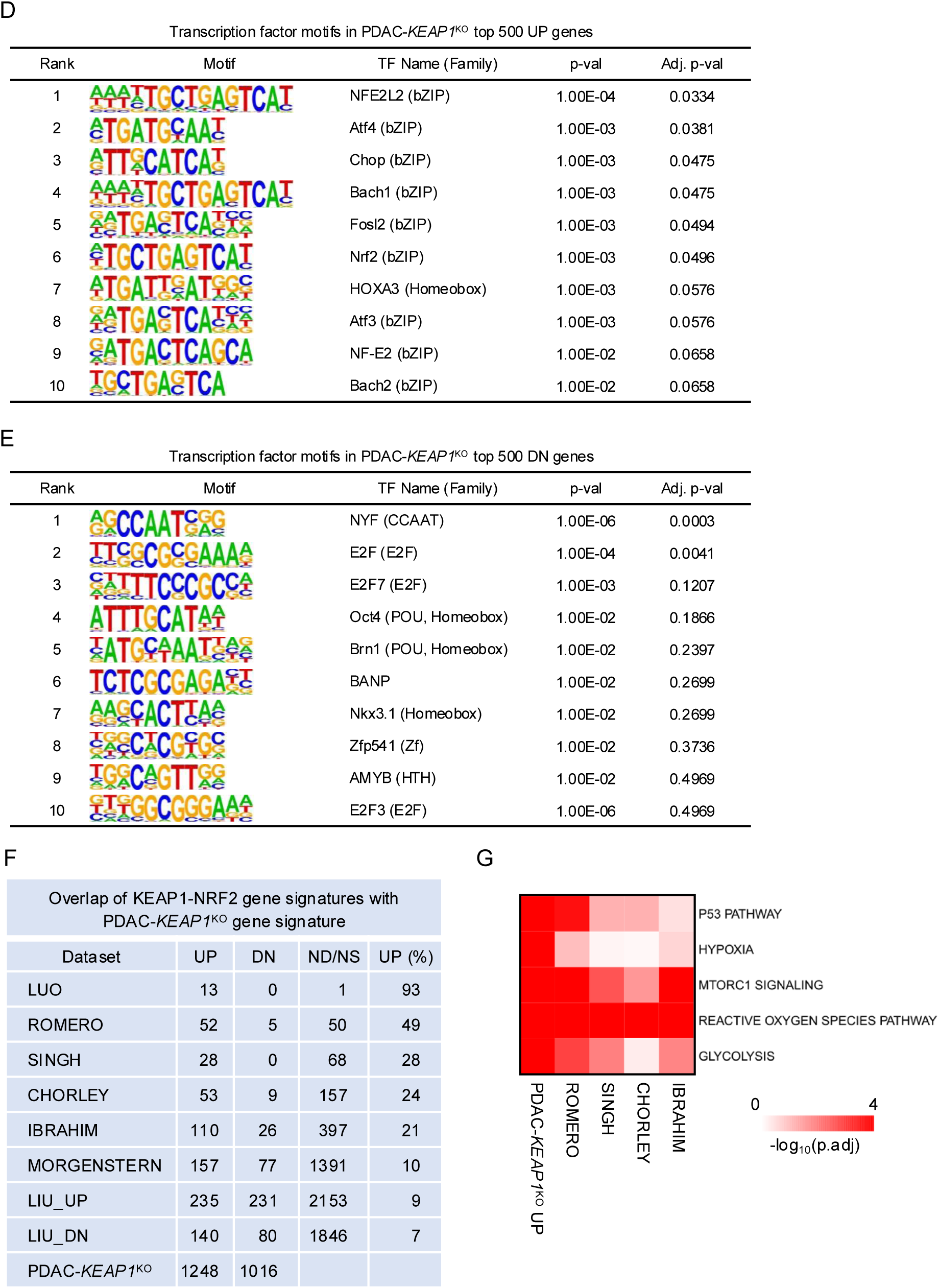
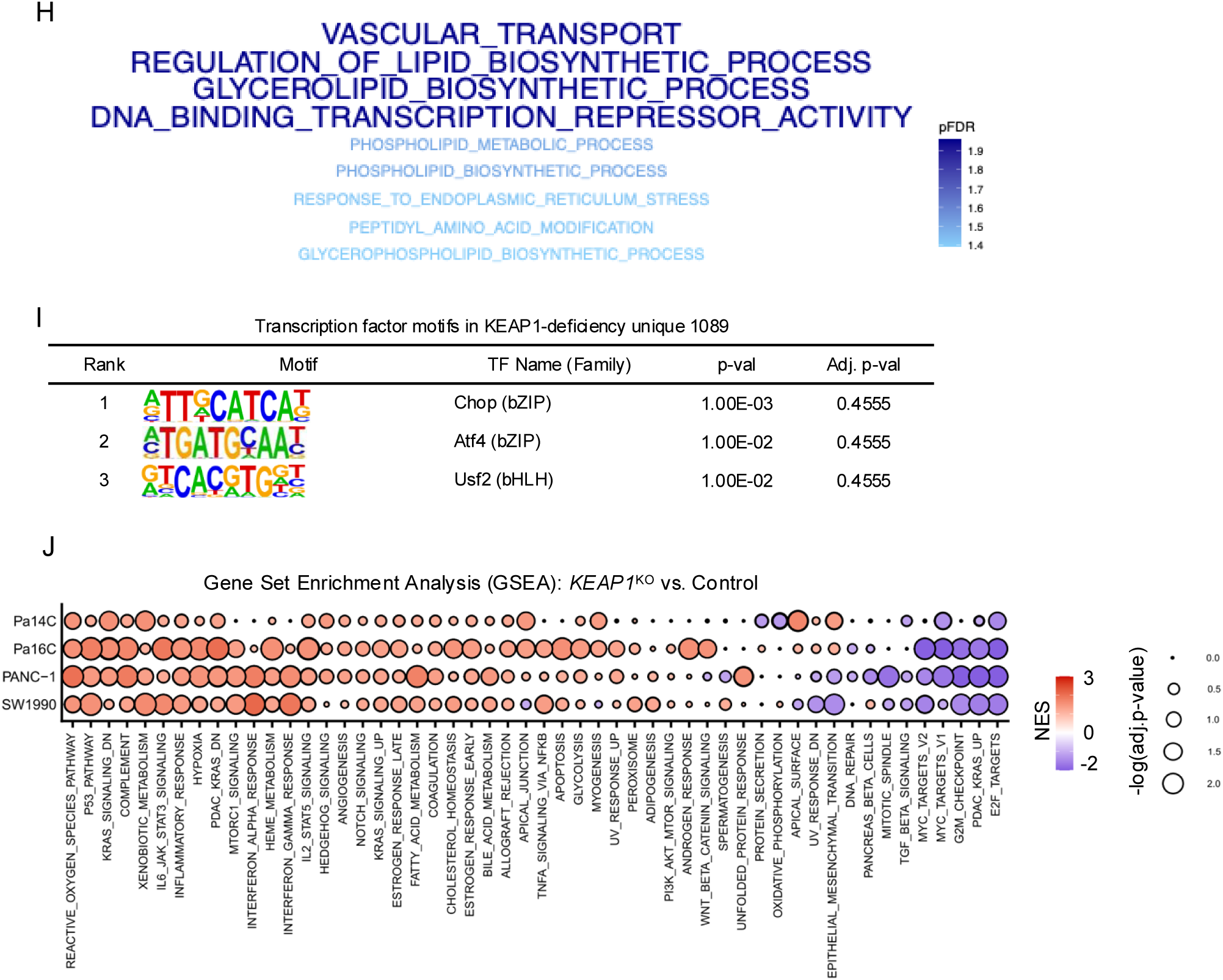
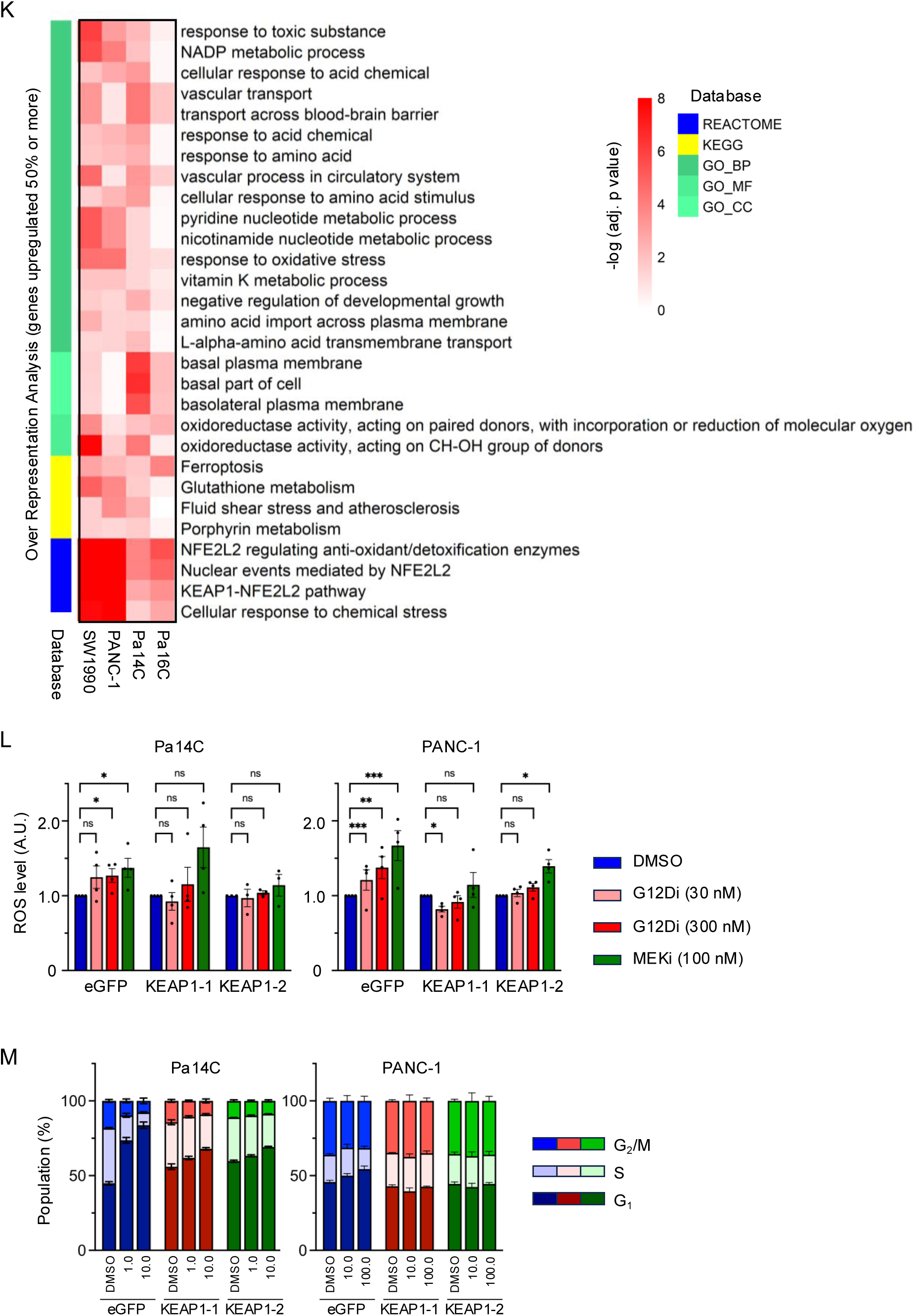
**A,** Representative western blots probing phospho-ERK 1/2 (pERK), total ERK 1/2 (ERK), phospho-RSK, total RSK, and vinculin in PANC-1 and Pa14C cells with knockout of *eGFP* or *KEAP1* (KEAP1-1, KEAP1-2) treated with RASi (0, 10, 100 nM) for 4 (top panel) or 24 hours (bottom panel). **B,** Representative western blots probing KEAP1, NRF2, SLC7A11, NQO1, and vinculin in PDAC cells (SW1990, Pa16C, Pa14C, PANC-1) induced for *KEAP1* knockout (+Dox), or non-induced controls (-Dox). **C,** PDAC cell lines (SW1990, Pa16C, Pa14C, PANC-1) induced for KEAP1 knockout (+Dox, red), or non-induced control (-Dox, blue), were treated with G12Di (0 – 300 nM) for 5 days, and cell viability was monitored by Calcein-AM assay at end of experiment. Normalized cell counts compared to the control conditions, which were normalized to 1.0, are plotted, and error bars denoted standard error of at least three independent biological replicates. **D,** Homer motif analysis was performed to identify the common transcription factor binding motifs for top 500 upregulated genes in *KEAP1*^KO^ PDAC cells. **E,** Homer motif analysis was performed to identify the common transcription factor binding motifs for top 500 downregulated genes in *KEAP1*^KO^ PDAC cells. **F,** The table summarizes genes within each signature and their differential expression trends in the PDAC-*KEAP1*^KO^ transcriptome. “UP” and “DN” denote significantly upregulated (log_2_FC >0, adj. p < 0.05) and downregulated (log_2_FC < 0, adj. p < 0.05) genes, respectively. “ND/NS” represents genes that were either not detected or non-significant (adj. p > 0.05). “UP (%)” indicates the percentage of upregulated genes within each signature in the PDAC-*KEAP1*^KO^ transcriptome. **G,** Top enriched pathways identified by ORA with 50 Hallmark gene sets on the PDAC *KEAP1*^KO^ UP, ROMERO, SINGH, CHORLEY, and IBRAHIM gene signatures**. H,** Top enriched pathways identified by ORA performed on 1,089 genes uniquely upregulated in the PDAC-*KEAP1*^KO^ transcriptome. Enrichment was determined using GO, KEGG, and Reactome databases. **I,** Homer motif analysis was performed to identify the common transcription factor binding motifs for 1,089 genes uniquely upregulated in *KEAP1*^KO^ PDAC cells. **J,** Gene Set Enrichment Analysis (GSEA) performed on *KEAP1*^KO^ PDAC cells using 50 Hallmark gene sets, as well as PDAC-KRAS UP and DN signatures previously reported by our group. **K,** ORA on genes with expression increased 50% or higher in PDAC cells identified commonly upregulated pathways from GO (biological process, BP; molecular function, MF; cellular component, CC), KEGG, and Reactome databases. **L,** Oxidative stress levels were monitored in *KEAP1*^KO^or control *eGFP*^KO^ Pa14C and PANC-1 cells treated with DMSO, G12Di (30 or 300 nM) or MEKi (100 nM) for 3 days. The ROS levels were determined using CellROX Green at the end of the experiment and read out using a Cytoflex S flow cytometry. **M,** Cell cycle population of Pa14C and PANC-1 with *KEAP1*^KO^or control *eGFP*^KO^ treated with indicated concentrations of G12Di or DMSO for 72 hours. Cells were stained using FxCycle™ PI/RNase Staining Solution and analyzed by a Cytoflex S flow cytometry.

**Supplementary Figure S4.**
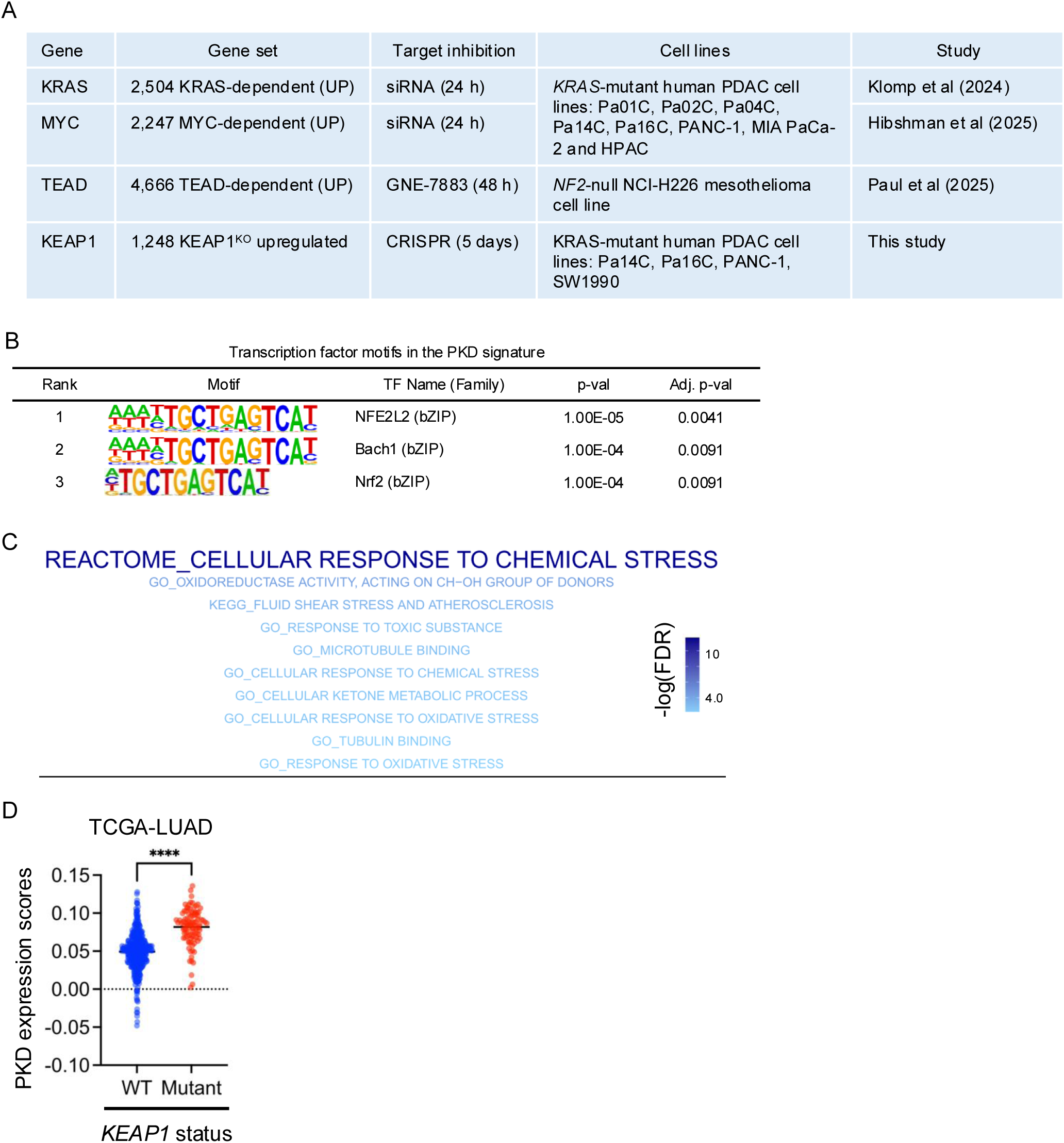
**A,** Table methodology and references of KRAS-, MYC-, and TEAD-dependent gene sets, as well as the *KEAP1*^KO^ upregulated gene set. **B,** Homer motif analysis was performed to identify the common transcription factor binding motifs for genes within the PKD signature. **C,** Top enriched pathways identified by ORA performed on the 200-gene PKD signature. Enrichment was determined with GO, KEGG, and Reactome databases. **D,** Gene expression scores of the PKD signature across TCGA-LUAD RNA expression datasets. Patients were categorized by their KEAP1 mutational status.

**Supplementary Figure S5.**
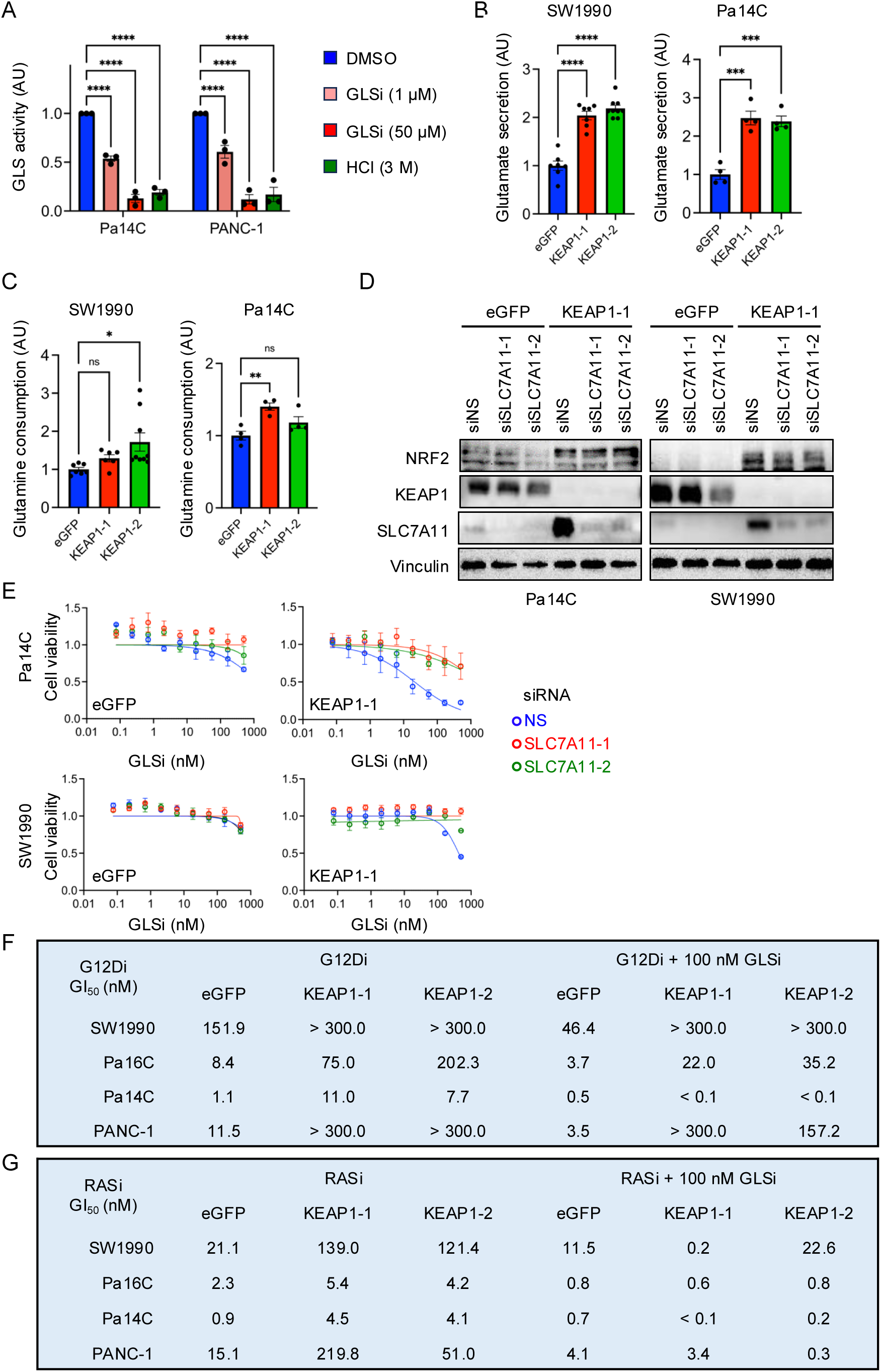
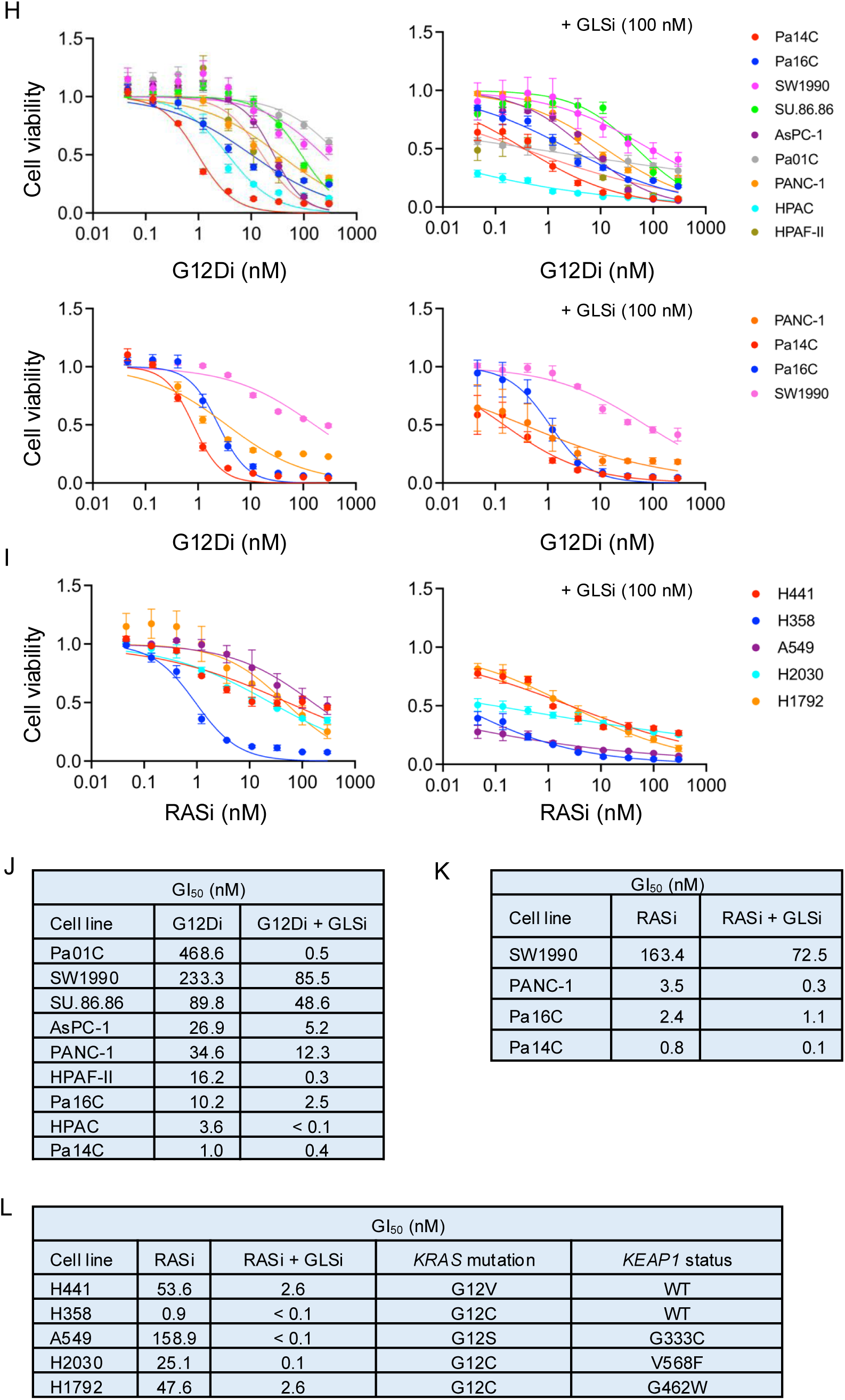
**A,** The glutaminase activity assay validated that GLSi is able to suppress mitochondrial glutaminase activity. High concentration (3 M) HCl was used as a negative control to quench all enzymatic reactions. **B,** Glutamate secretion was determined on conditioned medium collected from Pa14C and SW1990 cells with *KEAP1*^KO^ (KEAP1-1, red; and KEAP1-2, green) or *eGFP*^KO^ control cells by NMR. The error bars denoted standard errors of at least three independent biological replicates. **C,** Glutamine consumption was determined on conditioned medium collected from Pa14C and SW1990 cells with *KEAP1*^KO^ (KEAP1-1, red; and KEAP1-2, green) or *eGFP*^KO^ control cells by NMR. The error bars denoted standard errors of at least three independent biological replicates. **D,** Representative western blots probing NRF2, KEAP1, SLC7A11, and vinculin on *KEAP1*^KO^ (KEAP1-1) Pa14C and SW1990 cells, or their control counterparts (eGFP), treated with siRNA targeting SLC7A11 (siSLC7A11-1, siSLC7A11-2), or non-specific siRNA as controls (siNS). **E,** PDAC cell lines (Pa14C and SW1990) with *KEAP1*^KO^ knockout (KEAP-1), or *eGFP*^KO^ control cells, were treated with siRNAs targeting SLC7A11 (siSLC7A11-1, red; and siSLC7A11-2, green), or non-specific siRNA as a control (siNS, blue), and GLSi (0 – 500 nM) for 5 days, and cell viability was determined by the addition of Calcein-AM for 15 minutes at end of experiment. Normalized cell counts compared to the control conditions, which were normalized to 1.0, are plotted, and error bars denoted standard error of at least three independent biological replicates. **F,** Table of GI_50_ concentrations for *KEAP1*^KO^ or *eGFP*^KO^ PDAC cells treated with G12Di and with or without 100 nM GLSi. **G,** Table of GI_50_ concentrations for *KEAP1*^KO^ or *eGFP*^KO^ PDAC cells treated with RASi and with or without 100 nM GLSi. **H,** Parental PDAC cell lines treated with G12Di (0 – 300 nM) or RASi (0 – 300 nM) and with or without 100 nM GLSi for 5 days, and cell viability was determined by the addition of Calcein-AM for 15 minutes at end of experiment. Normalized cell counts compared to the control conditions, which were normalized to 1.0, are plotted, and error bars denoted standard error of at least three independent biological replicates. **I,** Parental LUAD cell lines treated with RASi (0 – 300 nM) and with or without 100 nM GLSi for 5 days, and cell viability was determined by the addition of Calcein-AM for 15 minutes at end of experiment. Normalized cell counts compared to the control conditions, which were normalized to 1.0, are plotted, and error bars denoted standard error of at least three independent biological replicates. **J,** Table of GI_50_ concentrations for parental PDAC cells treated with G12Di and with or without 100 nM GLSi. **K,** Table of GI_50_ concentrations for parental PDAC cells treated with RASi and with or without 100 nM GLSi. **L,** Table of GI_50_ concentrations for parental LUAD cells treated with RASi and with or without 100 nM GLSi.

**Supplementary Figure S6.**
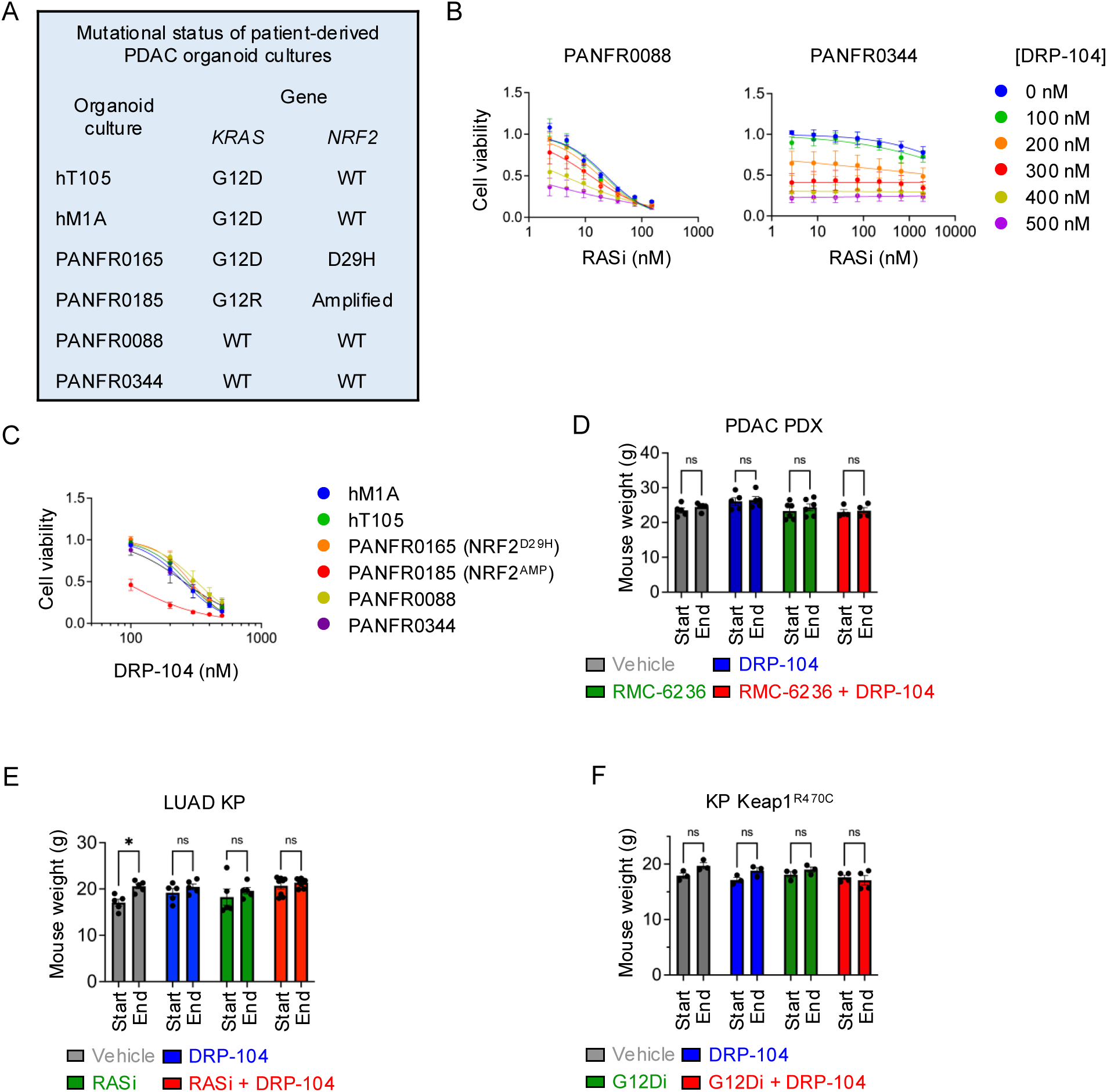
**A,** Table of *KRAS* and *NFE2L2/NRF2* mutational status of PDAC patient-derived organoids. **B,** PDAC patient derived organoids (PANFR0088, and PANFR0344) expressing wild type KRAS were treated with indicated concentrations of RASi (0 – 300 nM) or DRP-104 (0 – 500 nM) for 5 days, and cell viability was determined by 3D CellTiter Glo. Normalized cell viability compared to the control conditions, which were normalized to 1.0, were plotted, and error bars denoted standard error of at least three independent biological replicates. **C,** PDAC patient derived organoids treated with DRP-104 (0 – 500 nM) for 5 days, and cell viability was determined by 3D CellTiter Glo. Normalized cell viability compared to the control conditions, which were normalized to 1.0, were plotted, and error bars denoted standard error of at least three independent biological replicates. Data re-plotted from Fig. 6B and Supplementary Fig. S6B for visualizing DRP-104 single agent anti-tumor efficacy. **D,** Comparison of mouse body weights at the start and completion of the experiment from Figure 6C. **E,** Comparison of mouse body weights at the start and completion of the experiment from Figure 6D. **F,** Comparison of mouse body weights at the start and completion of the experiment from Figure 6E.

## REFERENCES

1. Varghese AM, Perry MA, Chou JF, Nandakumar S, Muldoon D, Erakky A, et al. Clinicogenomic landscape of pancreatic adenocarcinoma identifies KRAS mutant dosage as prognostic of overall survival. Nat Med 2025;31(2):466–77 doi 10.1038/s41591-024-03362-3.

2. Judd J, Abdel Karim N, Khan H, Naqash AR, Baca Y, Xiu J, et al. Characterization of KRAS Mutation Subtypes in Non-small Cell Lung Cancer. Mol Cancer Ther 2021;20(12):2577–84 doi 10.1158/1535-7163.MCT-21-0201.

3. Cox AD, Der CJ. “Undruggable KRAS”: druggable after all. Genes Dev 2025;39(1-2):132–62 doi 10.1101/gad.352081.124.

4. Skoulidis F, Li BT, Dy GK, Price TJ, Falchook GS, Wolf J, et al. Sotorasib for Lung Cancers with KRAS p.G12C Mutation. N Engl J Med 2021;384(25):2371–81 doi 10.1056/NEJMoa2103695.

5. Janne PA, Riely GJ, Gadgeel SM, Heist RS, Ou SI, Pacheco JM, et al. Adagrasib in Non-Small-Cell Lung Cancer Harboring a KRAS(G12C) Mutation. N Engl J Med 2022;387(2):120–31 doi 10.1056/NEJMoa2204619.

6. Fakih MG, Salvatore L, Esaki T, Modest DP, Lopez-Bravo DP, Taieb J, et al. Sotorasib plus Panitumumab in Refractory Colorectal Cancer with Mutated KRAS G12C. N Engl J Med 2023;389(23):2125–39 doi 10.1056/NEJMoa2308795.

7. Yaeger R, Uboha NV, Pelster MS, Bekaii-Saab TS, Barve M, Saltzman J, et al. Efficacy and Safety of Adagrasib plus Cetuximab in Patients with KRASG12C-Mutated Metastatic Colorectal Cancer. Cancer Discov 2024;14(6):982–93 doi 10.1158/2159-8290.CD-24-0217.

8. Isermann T, Sers C, Der CJ, Papke B. KRAS inhibitors: resistance drivers and combinatorial strategies. Trends Cancer 2025;11(2):91–116 doi 10.1016/j.trecan.2024.11.009.

9. Hallin J, Bowcut V, Calinisan A, Briere DM, Hargis L, Engstrom LD, et al. Anti-tumor efficacy of a potent and selective non-covalent KRAS(G12D) inhibitor. Nat Med 2022;28(10):2171–82 doi 10.1038/s41591-022-02007-7.

10. Weller C, Burnett GL, Jiang L, Chakraborty S, Zhang D, Vita NA, et al. A neomorphic protein interface catalyzes covalent inhibition of RAS(G12D) aspartic acid in tumors. Science 2025;389(6758):eads0239 doi 10.1126/science.ads0239.

11. Wolpin BM, Wainberg ZA, Garrido-Laguna I, Manji GA, Spira AI, Azad NS, et al. Trial in progress: RASolute 302—A phase 3, multicenter, global, open-label, randomized study of daraxonrasib (RMC-6236), a RAS(ON) multi-selective inhibitor, versus standard of care chemotherapy in patients with previously treated metastatic pancreatic ductal adenocarcinoma (PDAC). Journal of Clinical Oncology 2025;43(16_suppl):TPS4230-TPS doi 10.1200/JCO.2025.43.16_suppl.TPS4230.

12. Wasko UN, Jiang J, Dalton TC, Curiel-Garcia A, Edwards AC, Wang Y, et al. Tumor-selective activity of RAS-GTP inhibition in pancreatic cancer. Nature 2024 doi 10.1038/s41586-024-07379-z.

13. Ebright RY, Dilly J, Shaw AT, Aguirre AJ. Response and Resistance to RAS Inhibition in Cancer. Cancer Discov 2025;15(7):1325–49 doi 10.1158/2159-8290.CD-25-0349.

14. Aronchik I, Kar S, Zhuang Y, Ahler E, Lai LP, Seshadri V, et al. Abstract A081: Resistance Mechanisms to Monotherapy RAS(ON) Multi-Selective Inhibitor Daraxonrasib (RMC-6236) in RAS Mutant PDAC Inform Therapeutic Combination Strategies. Molecular Cancer Therapeutics 2025;24(10_Supplement):A081-A doi 10.1158/1535-7163.Targ-25-a081.

15. Sang B, Ye LF, Hu F, Pourfarjam Y, Cuevas-Navarro A, Fan S, et al. Mechanisms of resistance to active state selective tri-complex RAS inhibitors. bioRxiv 2025:2025.04.24.649345 doi 10.1101/2025.04.24.649345.

16. Awad MM, Liu S, Rybkin, II, Arbour KC, Dilly J, Zhu VW, et al. Acquired Resistance to KRAS(G12C) Inhibition in Cancer. N Engl J Med 2021;384(25):2382–93 doi 10.1056/NEJMoa2105281.

17. Zhao Y, Murciano-Goroff YR, Xue JY, Ang A, Lucas J, Mai TT, et al. Diverse alterations associated with resistance to KRAS(G12C) inhibition. Nature 2021;599(7886):679–83 doi 10.1038/s41586-021-04065-2.

18. Amodio V, Yaeger R, Arcella P, Cancelliere C, Lamba S, Lorenzato A, et al. EGFR Blockade Reverts Resistance to KRAS(G12C) Inhibition in Colorectal Cancer. Cancer Discov 2020;10(8):1129–39 doi 10.1158/2159-8290.CD-20-0187.

19. Holderfield M, Lee BJ, Jiang J, Tomlinson A, Seamon KJ, Mira A, et al. Concurrent inhibition of oncogenic and wild-type RAS-GTP for cancer therapy. Nature 2024 doi 10.1038/s41586-024-07205-6.

20. Edwards AC, Stalnecker CA, Jean Morales A, Taylor KE, Klomp JE, Klomp JA, et al. TEAD Inhibition Overcomes YAP1/TAZ-Driven Primary and Acquired Resistance to KRASG12C Inhibitors. Cancer Res 2023;83(24):4112–29 doi 10.1158/0008-5472.CAN-23-2994.

21. Tammaccaro SL, Prigent P, Le Bail JC, Dos-Santos O, Dassencourt L, Eskandar M, et al. TEAD Inhibitors Sensitize KRAS(G12C) Inhibitors via Dual Cell Cycle Arrest in KRAS(G12C)-Mutant NSCLC. Pharmaceuticals (Basel) 2023;16(4) doi 10.3390/ph16040553.

22. Dilly J, Hoffman MT, Abbassi L, Li Z, Paradiso F, Parent BD, et al. Mechanisms of resistance to oncogenic KRAS inhibition in pancreatic cancer. Cancer Discov 2024 doi 10.1158/2159-8290.CD-24-0177.

23. Negrao MV, Araujo HA, Lamberti G, Cooper AJ, Akhave NS, Zhou T, et al. Comutations and KRASG12C Inhibitor Efficacy in Advanced NSCLC. Cancer Discov 2023;13(7):1556–71 doi 10.1158/2159-8290.CD-22-1420.

24. Baird L, Yamamoto M. The Molecular Mechanisms Regulating the KEAP1-NRF2 Pathway. Mol Cell Biol 2020;40(13) doi 10.1128/MCB.00099-20.

25. Taguchi K, Yamamoto M. The KEAP1-NRF2 System as a Molecular Target of Cancer Treatment. Cancers (Basel) 2020;13(1) doi 10.3390/cancers13010046.

26. Pillai R, Hayashi M, Zavitsanou AM, Papagiannakopoulos T. NRF2: KEAPing Tumors Protected. Cancer Discov 2022;12(3):625–43 doi 10.1158/2159-8290.CD-21-0922.

27. Hast BE, Cloer EW, Goldfarb D, Li H, Siesser PF, Yan F, et al. Cancer-derived mutations in KEAP1 impair NRF2 degradation but not ubiquitination. Cancer Res 2014;74(3):808–17 doi 10.1158/0008-5472.CAN-13-1655.

28. Aboulkassim T, Tian X, Liu Q, Qiu D, Hancock M, Wu JH, et al. A NRF2 inhibitor selectively sensitizes KEAP1 mutant tumor cells to cisplatin and gefitinib by restoring NRF2-inhibitory function of KEAP1 mutants. Cell Rep 2023;42(9):113104 doi 10.1016/j.celrep.2023.113104.

29. Mukhopadhyay S, Goswami D, Adiseshaiah PP, Burgan W, Yi M, Guerin TM, et al. Undermining Glutaminolysis Bolsters Chemotherapy While NRF2 Promotes Chemoresistance in KRAS-Driven Pancreatic Cancers. Cancer Res 2020;80(8):1630–43 doi 10.1158/0008-5472.CAN-19-1363.

30. Krall EB, Wang B, Munoz DM, Ilic N, Raghavan S, Niederst MJ, et al. KEAP1 loss modulates sensitivity to kinase targeted therapy in lung cancer. Elife 2017;6 doi 10.7554/eLife.18970.

31. Lister A, Nedjadi T, Kitteringham NR, Campbell F, Costello E, Lloyd B, et al. Nrf2 is overexpressed in pancreatic cancer: implications for cell proliferation and therapy. Mol Cancer 2011;10:37 doi 10.1186/1476-4598-10-37.

32. Abu-Alainin W, Gana T, Liloglou T, Olayanju A, Barrera LN, Ferguson R, et al. UHRF1 regulation of the Keap1-Nrf2 pathway in pancreatic cancer contributes to oncogenesis. J Pathol 2016;238(3):423–33 doi 10.1002/path.4665.

33. Purohit V, Wang L, Yang H, Li J, Ney GM, Gumkowski ER, et al. ATDC binds to KEAP1 to drive NRF2-mediated tumorigenesis and chemoresistance in pancreatic cancer. Genes Dev 2021;35(3-4):218–33 doi 10.1101/gad.344184.120.

34. DeNicola GM, Karreth FA, Humpton TJ, Gopinathan A, Wei C, Frese K, et al. Oncogene-induced Nrf2 transcription promotes ROS detoxification and tumorigenesis. Nature 2011;475(7354):106–9 doi 10.1038/nature10189.

35. Drosten M, Dhawahir A, Sum EY, Urosevic J, Lechuga CG, Esteban LM, et al. Genetic analysis of Ras signalling pathways in cell proliferation, migration and survival. EMBO J 2010;29(6):1091–104 doi 10.1038/emboj.2010.7.

36. Waters AM, Ozkan-Dagliyan I, Vaseva AV, Fer N, Strathern LA, Hobbs GA, et al. Evaluation of the selectivity and sensitivity of isoform- and mutation-specific RAS antibodies. Sci Signal 2017;10(498) doi 10.1126/scisignal.aao3332.

37. Lin KH, Rutter JC, Xie A, Pardieu B, Winn ET, Bello RD, et al. Using antagonistic pleiotropy to design a chemotherapy-induced evolutionary trap to target drug resistance in cancer. Nat Genet 2020;52(4):408–17 doi 10.1038/s41588-020-0590-9.

38. Sacher A, LoRusso P, Patel MR, Miller WH, Jr., Garralda E, Forster MD, et al. Single-Agent Divarasib (GDC-6036) in Solid Tumors with a KRAS G12C Mutation. N Engl J Med 2023;389(8):710–21 doi 10.1056/NEJMoa2303810.

39. Negrao MV, Paula AG, Molkentine D, Hover L, Nilsson M, Vokes N, et al. Impact of Co-mutations and Transcriptional Signatures in Non-Small Cell Lung Cancer Patients Treated with Adagrasib in the KRYSTAL-1 Trial. Clin Cancer Res 2025;31(6):1069–81 doi 10.1158/1078-0432.CCR-24-2310.

40. Hayes JD, Dinkova-Kostova AT. The Nrf2 regulatory network provides an interface between redox and intermediary metabolism. Trends Biochem Sci 2014;39(4):199–218 doi 10.1016/j.tibs.2014.02.002.

41. Hur W, Sun Z, Jiang T, Mason DE, Peters EC, Zhang DD, et al. A small-molecule inducer of the antioxidant response element. Chem Biol 2010;17(5):537–47 doi 10.1016/j.chembiol.2010.03.013.

42. Robledinos-Anton N, Fernandez-Gines R, Manda G, Cuadrado A. Activators and Inhibitors of NRF2: A Review of Their Potential for Clinical Development. Oxid Med Cell Longev 2019;2019:9372182 doi 10.1155/2019/9372182.

43. Romero R, Sayin VI, Davidson SM, Bauer MR, Singh SX, LeBoeuf SE, et al. Keap1 loss promotes Kras-driven lung cancer and results in dependence on glutaminolysis. Nat Med 2017;23(11):1362–8 doi 10.1038/nm.4407.

44. Zavitsanou AM, Pillai R, Hao Y, Wu WL, Bartnicki E, Karakousi T, et al. KEAP1 mutation in lung adenocarcinoma promotes immune evasion and immunotherapy resistance. Cell Rep 2023;42(11):113295 doi 10.1016/j.celrep.2023.113295.

45. Klomp JA, Klomp JE, Stalnecker CA, Bryant KL, Edwards AC, Drizyte-Miller K, et al. Defining the KRAS- and ERK-dependent transcriptome in KRAS-mutant cancers. Science 2024;384(6700):eadk0775 doi 10.1126/science.adk0775.

46. Klomp JE, Diehl JN, Klomp JA, Edwards AC, Yang R, Morales AJ, et al. Determining the ERK-regulated phosphoproteome driving KRAS-mutant cancer. Science 2024;384(6700):eadk0850 doi 10.1126/science.adk0850.

47. Barger CJ, Branick C, Chee L, Karpf AR. Pan-Cancer Analyses Reveal Genomic Features of FOXM1 Overexpression in Cancer. Cancers (Basel) 2019;11(2) doi 10.3390/cancers11020251.

48. Heinz S, Benner C, Spann N, Bertolino E, Lin YC, Laslo P, et al. Simple combinations of lineage-determining transcription factors prime cis-regulatory elements required for macrophage and B cell identities. Mol Cell 2010;38(4):576–89 doi 10.1016/j.molcel.2010.05.004.

49. Raghunath A, Sundarraj K, Nagarajan R, Arfuso F, Bian J, Kumar AP, et al. Antioxidant response elements: Discovery, classes, regulation and potential applications. Redox Biol 2018;17:297–314 doi 10.1016/j.redox.2018.05.002.

50. Chorley BN, Campbell MR, Wang X, Karaca M, Sambandan D, Bangura F, et al. Identification of novel NRF2-regulated genes by ChIP-Seq: influence on retinoid X receptor alpha. Nucleic Acids Res 2012;40(15):7416–29 doi 10.1093/nar/gks409.

51. Ibrahim L, Mesgarzadeh J, Xu I, Powers ET, Wiseman RL, Bollong MJ. Defining the Functional Targets of Cap’n’collar Transcription Factors NRF1, NRF2, and NRF3. Antioxidants (Basel) 2020;9(10) doi 10.3390/antiox9101025.

52. Liu P, Kerins MJ, Tian W, Neupane D, Zhang DD, Ooi A. Differential and overlapping targets of the transcriptional regulators NRF1, NRF2, and NRF3 in human cells. J Biol Chem 2019;294(48):18131–49 doi 10.1074/jbc.RA119.009591.

53. Luo G, Kumar H, Aldridge K, Rieger S, Han E, Jiang E, et al. A Core NRF2 Gene Set Defined Through Comprehensive Transcriptomic Analysis Predicts Selective Drug Resistance and Poor Multicancer Prognosis. Antioxid Redox Signal 2024;41(16-18):1031–50 doi 10.1089/ars.2023.0409.

54. Morgenstern C, Lastres-Becker I, Demirdogen BC, Costa VM, Daiber A, Foresti R, et al. Biomarkers of NRF2 signalling: Current status and future challenges. Redox Biol 2024;72:103134 doi 10.1016/j.redox.2024.103134.

55. Singh A, Daemen A, Nickles D, Jeon SM, Foreman O, Sudini K, et al. NRF2 Activation Promotes Aggressive Lung Cancer and Associates with Poor Clinical Outcomes. Clin Cancer Res 2021;27(3):877–88 doi 10.1158/1078-0432.CCR-20-1985.

56. Sayin VI, LeBoeuf SE, Singh SX, Davidson SM, Biancur D, Guzelhan BS, et al. Activation of the NRF2 antioxidant program generates an imbalance in central carbon metabolism in cancer. Elife 2017;6 doi 10.7554/eLife.28083.

57. Liberzon A, Birger C, Thorvaldsdottir H, Ghandi M, Mesirov JP, Tamayo P. The Molecular Signatures Database (MSigDB) hallmark gene set collection. Cell Syst 2015;1(6):417–25 doi 10.1016/j.cels.2015.12.004.

58. DeLiberty JM, Roach MK, Stalnecker CA, Robb R, Schechter EG, Pieper NL, et al. Concurrent Inhibition of the RAS-MAPK Pathway and PIKfyve is a Therapeutic Strategy for Pancreatic Cancer. Cancer Res 2025 doi 10.1158/0008-5472.CAN-24-1757.

59. Mohanty A, Nam A, Srivastava S, Jones J, Lomenick B, Singhal SS, et al. Acquired resistance to KRAS G12C small-molecule inhibitors via genetic/nongenetic mechanisms in lung cancer. Sci Adv 2023;9(41):eade3816 doi 10.1126/sciadv.ade3816.

60. Lastra D, Escoll M, Cuadrado A. Transcription Factor NRF2 Participates in Cell Cycle Progression at the Level of G1/S and Mitotic Checkpoints. Antioxidants (Basel) 2022;11(5) doi 10.3390/antiox11050946.

61. Reddy NM, Kleeberger SR, Bream JH, Fallon PG, Kensler TW, Yamamoto M, et al. Genetic disruption of the Nrf2 compromises cell-cycle progression by impairing GSH-induced redox signaling. Oncogene 2008;27(44):5821–32 doi 10.1038/onc.2008.188.

62. Hibshman PS, Stalnecker CA, Klomp JA, Drizyte-Miller K, Klomp JE, Edwards AC, et al. Defining the MYC-regulated transcriptome and kinome that support KRAS- and ERK-dependent growth of pancreatic cancer. Sci Signal 2025;18(906):eadu7145 doi 10.1126/scisignal.adu7145.

63. Paul S, Hagenbeek TJ, Tremblay J, Kameswaran V, Ong C, Liu C, et al. Cooperation between the Hippo and MAPK pathway activation drives acquired resistance to TEAD inhibition. Nat Commun 2025;16(1):1743 doi 10.1038/s41467-025-56634-y.

64. Luo Y, Hitz BC, Gabdank I, Hilton JA, Kagda MS, Lam B, et al. New developments on the Encyclopedia of DNA Elements (ENCODE) data portal. Nucleic Acids Res 2020;48(D1):D882–D9 doi 10.1093/nar/gkz1062.

65. Dhakshinamoorthy S, Jain AK, Bloom DA, Jaiswal AK. Bach1 competes with Nrf2 leading to negative regulation of the antioxidant response element (ARE)-mediated NAD(P)H:quinone oxidoreductase 1 gene expression and induction in response to antioxidants. J Biol Chem 2005;280(17):16891–900 doi 10.1074/jbc.M500166200.

66. Balaji SA, Udupa N, Chamallamudi MR, Gupta V, Rangarajan A. Role of the Drug Transporter ABCC3 in Breast Cancer Chemoresistance. PLoS One 2016;11(5):e0155013 doi 10.1371/journal.pone.0155013.

67. Girardi E, Cesar-Razquin A, Lindinger S, Papakostas K, Konecka J, Hemmerich J, et al. A widespread role for SLC transmembrane transporters in resistance to cytotoxic drugs. Nat Chem Biol 2020;16(4):469–78 doi 10.1038/s41589-020-0483-3.

68. Nigam SK. The SLC22 Transporter Family: A Paradigm for the Impact of Drug Transporters on Metabolic Pathways, Signaling, and Disease. Annu Rev Pharmacol Toxicol 2018;58:663–87 doi 10.1146/annurev-pharmtox-010617-052713.

69. Penning TM, Jonnalagadda S, Trippier PC, Rizner TL. Aldo-Keto Reductases and Cancer Drug Resistance. Pharmacol Rev 2021;73(3):1150–71 doi 10.1124/pharmrev.120.000122.

70. Matsunaga T, Hojo A, Yamane Y, Endo S, El-Kabbani O, Hara A. Pathophysiological roles of aldo-keto reductases (AKR1C1 and AKR1C3) in development of cisplatin resistance in human colon cancers. Chem Biol Interact 2013;202(1-3):234–42 doi 10.1016/j.cbi.2012.09.024.

71. Matsunaga T, Wada Y, Endo S, Soda M, El-Kabbani O, Hara A. Aldo-Keto Reductase 1B10 and Its Role in Proliferation Capacity of Drug-Resistant Cancers. Front Pharmacol 2012;3:5 doi 10.3389/fphar.2012.00005.

72. Penning TM. Aldo-Keto Reductase Regulation by the Nrf2 System: Implications for Stress Response, Chemotherapy Drug Resistance, and Carcinogenesis. Chem Res Toxicol 2017;30(1):162–76 doi 10.1021/acs.chemrestox.6b00319.

73. Bryant KL, Stalnecker CA, Zeitouni D, Klomp JE, Peng S, Tikunov AP, et al. Combination of ERK and autophagy inhibition as a treatment approach for pancreatic cancer. Nat Med 2019;25(4):628–40 doi 10.1038/s41591-019-0368-8.

74. Kinsey CG, Camolotto SA, Boespflug AM, Guillen KP, Foth M, Truong A, et al. Protective autophagy elicited by RAF-->MEK-->ERK inhibition suggests a treatment strategy for RAS-driven cancers. Nat Med 2019;25(4):620–7 doi 10.1038/s41591-019-0367-9.

75. Waters AM, Khatib TO, Papke B, Goodwin CM, Hobbs GA, Diehl JN, et al. Targeting p130Cas- and microtubule-dependent MYC regulation sensitizes pancreatic cancer to ERK MAPK inhibition. Cell Rep 2021;35(13):109291 doi 10.1016/j.celrep.2021.109291.

76. Zhang S, Huang WC, Li P, Guo H, Poh SB, Brady SW, et al. Combating trastuzumab resistance by targeting SRC, a common node downstream of multiple resistance pathways. Nat Med 2011;17(4):461–9 doi 10.1038/nm.2309.

77. Duxbury MS, Ito H, Zinner MJ, Ashley SW, Whang EE. Inhibition of SRC tyrosine kinase impairs inherent and acquired gemcitabine resistance in human pancreatic adenocarcinoma cells. Clin Cancer Res 2004;10(7):2307–18 doi 10.1158/1078-0432.ccr-1183-3.

78. Salmon M, Alvarez-Diaz R, Fustero-Torre C, Brehey O, Lechuga CG, Sanclemente M, et al. Kras oncogene ablation prevents resistance in advanced lung adenocarcinomas. J Clin Invest 2023;133(7) doi 10.1172/JCI164413.

79. Yang WH, Qiu Y, Stamatatos O, Janowitz T, Lukey MJ. Enhancing the Efficacy of Glutamine Metabolism Inhibitors in Cancer Therapy. Trends Cancer 2021;7(8):790–804 doi 10.1016/j.trecan.2021.04.003.

80. Nguyen TT, Katt WP, Cerione RA. Alone and together: current approaches to targeting glutaminase enzymes as part of anti-cancer therapies. Future Drug Discov 2023;4(4):FDD79 doi 10.4155/fdd-2022-0011.

81. Suzuki T, Muramatsu A, Saito R, Iso T, Shibata T, Kuwata K, et al. Molecular Mechanism of Cellular Oxidative Stress Sensing by Keap1. Cell Rep 2019;28(3):746–58 e4 doi 10.1016/j.celrep.2019.06.047.

82. Badgley MA, Kremer DM, Maurer HC, DelGiorno KE, Lee HJ, Purohit V, et al. Cysteine depletion induces pancreatic tumor ferroptosis in mice. Science 2020;368(6486):85–9 doi 10.1126/science.aaw9872.

83. Koppula P, Zhuang L, Gan B. Cystine transporter SLC7A11/xCT in cancer: ferroptosis, nutrient dependency, and cancer therapy. Protein Cell 2021;12(8):599–620 doi 10.1007/s13238-020-00789-5.

84. Biancur DE, Paulo JA, Malachowska B, Quiles Del Rey M, Sousa CM, Wang X, et al. Compensatory metabolic networks in pancreatic cancers upon perturbation of glutamine metabolism. Nat Commun 2017;8:15965 doi 10.1038/ncomms15965.

85. Rais R, Lemberg KM, Tenora L, Arwood ML, Pal A, Alt J, et al. Discovery of DRP-104, a tumor-targeted metabolic inhibitor prodrug. Sci Adv 2022;8(46):eabq5925 doi 10.1126/sciadv.abq5925.

86. Pillai R, LeBoeuf SE, Hao Y, New C, Blum JLE, Rashidfarrokhi A, et al. Glutamine antagonist DRP-104 suppresses tumor growth and enhances response to checkpoint blockade in KEAP1 mutant lung cancer. Sci Adv 2024;10(13):eadm9859 doi 10.1126/sciadv.adm9859.

87. Encarnacion-Rosado J, Sohn ASW, Biancur DE, Lin EY, Osorio-Vasquez V, Rodrick T, et al. Targeting pancreatic cancer metabolic dependencies through glutamine antagonism. Nat Cancer 2024;5(1):85–99 doi 10.1038/s43018-023-00647-3.

88. Fukutomi T, Takagi K, Mizushima T, Ohuchi N, Yamamoto M. Kinetic, thermodynamic, and structural characterizations of the association between Nrf2-DLGex degron and Keap1. Mol Cell Biol 2014;34(5):832–46 doi 10.1128/MCB.01191-13.

89. Jiang J, Jiang L, Maldonato BJ, Wang Y, Holderfield M, Aronchik I, et al. Translational and Therapeutic Evaluation of RAS-GTP Inhibition by RMC-6236 in RAS-Driven Cancers. Cancer Discov 2024:OF1-OF24 doi 10.1158/2159-8290.CD-24-0027.

90. Dhanasekaran R, Deutzmann A, Mahauad-Fernandez WD, Hansen AS, Gouw AM, Felsher DW. The MYC oncogene - the grand orchestrator of cancer growth and immune evasion. Nat Rev Clin Oncol 2022;19(1):23–36 doi 10.1038/s41571-021-00549-2.

91. Cunningham R, Hansen CG. The Hippo pathway in cancer: YAP/TAZ and TEAD as therapeutic targets in cancer. Clin Sci (Lond) 2022;136(3):197–222 doi 10.1042/CS20201474.

92. Boj SF, Hwang CI, Baker LA, Chio, II, Engle DD, Corbo V, et al. Organoid models of human and mouse ductal pancreatic cancer. Cell 2015;160(1-2):324–38 doi 10.1016/j.cell.2014.12.021.

93. Li W, Xu H, Xiao T, Cong L, Love MI, Zhang F, et al. MAGeCK enables robust identification of essential genes from genome-scale CRISPR/Cas9 knockout screens. Genome Biol 2014;15(12):554 doi 10.1186/s13059-014-0554-4.

94. Stalnecker CA, Ulrich SM, Li Y, Ramachandran S, McBrayer MK, DeBerardinis RJ, et al. Mechanism by which a recently discovered allosteric inhibitor blocks glutamine metabolism in transformed cells. Proc Natl Acad Sci U S A 2015;112(2):394–9 doi 10.1073/pnas.1414056112.

95. Nguyen TT, Ramachandran S, Hill MJ, Cerione RA. High-resolution structures of mitochondrial glutaminase C tetramers indicate conformational changes upon phosphate binding. J Biol Chem 2022;298(2):101564 doi 10.1016/j.jbc.2022.101564.

96. Foroutan M, Bhuva DD, Lyu R, Horan K, Cursons J, Davis MJ. Single sample scoring of molecular phenotypes. BMC Bioinformatics 2018;19(1):404 doi 10.1186/s12859-018-2435-4.

97. Frankish A, Diekhans M, Ferreira AM, Johnson R, Jungreis I, Loveland J, et al. GENCODE reference annotation for the human and mouse genomes. Nucleic Acids Res 2019;47(D1):D766–D73 doi 10.1093/nar/gky955.

98. Lawrence M, Huber W, Pages H, Aboyoun P, Carlson M, Gentleman R, et al. Software for computing and annotating genomic ranges. PLoS Comput Biol 2013;9(8):e1003118 doi 10.1371/journal.pcbi.1003118.

99. Durinck S, Moreau Y, Kasprzyk A, Davis S, De Moor B, Brazma A, et al. BioMart and Bioconductor: a powerful link between biological databases and microarray data analysis. Bioinformatics 2005;21(16):3439–40 doi 10.1093/bioinformatics/bti525.

