## Supplementary Table S2 for "Characterization and therapeutic suppression of KEAP1-NRF2-driven resistance to KRAS inhibitors in pancreatic and lung cancer"

| Table S1. KEAP1-NRF2 gene signatures | | | |
| --- | --- | --- | --- |
| Gene set | Size | Species | Description |
| Chorley | 232 | H | Microarray analyses of six human lymphoblastoid cells treated with SFN (NRF2 inducer; 5 h) (291 UP and 217 DN genes; 508 total genes) and NRF2 ChIP-seq (849 binding sites); identified 67 SFN-responsive genes with ChIP-seq binding (Chorley et al, 2010) |
| IBRAHIM | 533 | H | RNA-Seq analyses HEK293T SV40 large T antigen expressing human embryonic epithelial cells transiently (2 h) overexpressing a KEAP1 binding-deficient NRF2^T80D^ mutant identified 2,141 UP and 1,388 DN genes); overlap with Liu et al, 2019 (2,619 UP and 2,067 DN genes) allowed identification of a consensus gene set of 239 high confidence NRF2-regulated genes (Ibrahim et al, 2020) |
| LIU | 167 | H | Dox-inducible FLAG-tagged NRF2 in human U2OS osteosarcoma cell line and sulforaphane to facilitate NRF2 translocation and RNA-Seq identified 2,619 UP and 2,067 DN genes; combined with ChIP data, 167 genes shared by NRF1-3 (Liu et al, 2019) |
| LUO | 14 | H | Generated from 7 transcriptomic databases: treatment of RPMI-8226 myeloma, OVCAR-8 ovarian, SF8286 glioma and primary dermal fibroblasts with NRF2 activator CDDO-2P-Im (6 h), KEAP1 CRISPR in ARH-77 B cell lymphoblastic, KEAP1/NRF2 mutated versus WT NSCLC in TCGA and NSCLC cell lines in CCLE; defined a core set of 14 NRF2 UP genes (Luo et al, 2024) |
| Morgenstern | 1,625 | M, R, H | Literature search (238 mouse, rat or human studies) for (1) ChIP direct targets, (2) NRF2 suppression DN genes, or (3) NRF2 activation UP genes identified 1,625 NRF2-stimulated genes/proteins in at least one category, and 6 that met all 3 criteria (Morgenstern et al, 2024) |
| SINGH | 96 | H | Differential analysis using RNA-seq from TCGA dataset comparing KEAP1 or NRF2 altered vs. KEAP1 and NRF2 WT in 439 human lung adenocarcinoma tumors (Singh et al, 2021) |
| ROMERO | 108 | M, H | Common genes from previously established KEAP1-NRF2 gene sets from Singh et al, 2008; Mitsuishi et al, 2012; and Malhotra et al, 2010 (Romero et al, 2017) |
